# Age-related changes in MEG theta and alpha oscillatory signatures of attentional control

**DOI:** 10.1101/461020

**Authors:** Eleanor Huizeling, Hongfang Wang, Carol Holland, Klaus Kessler

## Abstract

In our recent behavioural research (Callaghan et al., 2017), we reported age-related changes in the costs of switching between from a temporal (rapid serial visual presentation) to a spatial (visual search) attention task. Using magnetoencephalography, we have now compared the neural signatures of attention refocusing between three age groups (19-30, 40-49 and 60+ years) and found differences in task-related modulation and cortical localisation of alpha and theta oscillations. Efficient, faster switching between the temporal and spatial attention tasks in the youngest group compared to both older groups was reflected in parietal theta effects that were significantly reduced in the older groups. Residual parietal theta activity in older individuals was, however, beneficial to attentional switching, and could reflect the preservation of attention mechanisms. Difficulties in refocusing attention in the older and middle-aged adults (slowed response times) were accompanied by reduced theta power modulation in occipital and cerebellar regions. In addition to this posterior theta deficit, older and middle-aged adults presented with increased recruitment of frontal (both groups) and temporal (older group) areas. Theta oscillations therefore demonstrated a posterior to anterior processing shift, which may reflect the engagement of alternative strategies in older- and middle-age, such as top-down attentional control in an attempt to compensate for posterior processing deficits. Additional frontal theta recruitment indeed appeared to be beneficial for faster performance, as reflected by correlational analysis. Temporal theta (in older-age), however, did not correlate with performance, but could reflect strategies unrelated to speeded performance (e.g. silent vocalisation to maintain task goals). Alpha oscillations did not significantly correlate with task performance, possibly reflecting decreased neural precision or de-differentiation, but require further investigation.

## Introduction

Over past decades the predominant view of age-related changes in cognitive function was that of a continuous decline (Dempster, 1992; Salthouse, 1996; West, 1996). However, recent findings suggest that older adults are able to efficiently recruit alternative cognitive mechanisms when performing cognitive tasks (Cabeza et al., 2018; Daselaar et al., 2015; Park and Reuter-Lorenz, 2009; Reuter-Lorenz and Park, 2014). An important trajectory of research has emerged that aims to find out which areas of cognition remain high-functioning for longer and can be utilised to support lesser preserved processes. Here we aimed to understand how flexible refocusing of attention in time and space might be affected in middle- and older age and whether certain processing elements, such as bottom-up stimulus-driven processing, are affected more strongly than others, such as top-down attentional control (or vice versa). A further aim was to investigate whether preserved functioning could be recruited to support more affected processing elements.

Age-related deterioration of performance has been reported separately for temporal as well as spatial selective attention (Bennett et al., 2012; Foster et al., 1995; Humphrey and Kramer, 1997; Lahar et al., 2001; Lee and Hsieh, 2009; Maciokas and Crognale, 2003; Nagamatsu et al., 2013; Plude and Doussard-Roosevelt, 1989). However, abilities in switching attention from one mode to the other have remained under-investigated (Callaghan et al., 2017), despite dynamic refocusing of attention potentially being crucial for everyday activities such as driving (Callaghan et al., 2017; Huizeling et al., 2020; Torrens-Burton et al., 2020). The aim of the current study was to investigate the neural patterns that reflect age-related changes in the ability to refocus or reallocate attention between time and space. Using a paradigm developed in our recent behavioural work (Callaghan et al., 2017), we compared three age groups on their ability to switch from a standard temporal attention task, requiring identification of a single target in a stream of distractors (rapid serial visual presentation, RSVP), to allocating attention spatially, to identify a target in a visual search (VS) task.

Spatial attention is typically quantified using VS tasks, where participants’ response times (RTs) to detect a predefined visual target among an array of distractors are recorded (Treisman, 1985; Treisman and Gormican, 1988). The target can either “pop-out” from the distractors by being different on a single, salient dimension (e.g. by being the only “T” among “O” distractors) or the target may share more than one feature dimension with distractors (e.g. a red “T” among red “O” and blue “T” distractors). The former is termed “pop-out” VS as RTs do not depend on the number of distractors, while the latter is termed serial VS as RTs depend on the number of distractors and VS is assumed to rely on a sequential scanning process of all stimuli until the target is found. Serial VS but not pop-out VS has been reported to reveal age-related decline in addition to a basic general slowing of processing speed, indicating possible deficiencies in the sequential engaging and disengaging of attention during serial VS (Bennett et al., 2012; Foster et al., 1995; Humphrey and Kramer, 1997; Nagamatsu et al., 2013; Plude and Doussard-Roosevelt, 1989). However, when Callaghan et al. (2017) presented VS tasks preceded by temporal attention tasks (single-target RSVP), attentional demands were increased and ceiling effects during serial search appeared to prevent increased age-related costs to clearly emerge. In contrast, pop-out search was faster than serial search and revealed age-related increases in RTs compatible with enhanced difficulties of switching from a temporal to a spatial focus of attention (Callaghan et al., 2017). In the current study we therefore employed pop-out VS only, presented in succession to a single-target RSVP task.

Age-related declines in temporal attention have also been reported, which is typically tested using variations of RSVP streams of stimuli (e.g. letters or digits). Older adults are not only slower at processing visual stimuli (Ball et al., 2006; Rubin et al., 2007) but also display an increased magnitude of the “attentional blink” effect. The attentional blink effect is manifest when, for up to 500ms after detecting a (first) target in an RSVP stream there is a reduced ability to detect a second target (Raymond et al., 1992). This effect is stronger and lasts for longer with increased age (Lahar et al., 2001; Lee and Hsieh, 2009; Maciokas and Crognale, 2003; Shih, 2009; van Leeuwen et al., 2009), which, again, cannot be explained by general slowing alone (Lee and Hsieh, 2009; Maciokas and Crognale, 2003). In a recent behavioural study (Callaghan et al., 2017), we investigated whether further costs are incurred with age when selective attention mechanisms have to be re-tuned or switched from selectively attending to a target in an RSVP stream to selecting a target in a VS display. We indeed observed increased “Switch-Costs” in older age groups, especially during pop-out VS. The current study set out to investigate the age-related neural patterns that might underlie this reduced attentional flexibility.

Overlapping brain networks across occipital, frontal, parietal and motor regions have been implicated in directing attention in both time and space (Coull and Nobre, 1998; Fu et al., 2005; Gross et al., 2004; Madden et al., 2007; Nagamatsu et al., 2013; Shapiro et al., 2002).

In addition to finding overlapping activation for temporal and spatial attention in their functional magnetic resonance imaging (fMRI) and positron emission tomography (PET) studies, Coull and Nobre (1998) also found sub-patterns of activation that were distinct for the two types of attention. The latter suggests that the human brain might have to be “re-tuned” when switching from a temporal to a spatial focus of attention (and vice versa); a dynamic process that could be particularly affected by age-related decline. For our current study we therefore expected fronto-parietal networks in conjunction with occipital areas to reveal age-related changes. To complicate matters, findings are somewhat inconsistent in terms of whether reduced activity in these cortical attention networks (Cabeza, 2002; Madden and Gottlob, 1997; Madden et al., 2002; Ross et al., 1997) or more widely distributed activity across the cortex (Adamo et al., 2003; Lague-Beauvais et al., 2013; Madden et al., 2007; Nagamatsu et al., 2013) is the primary reflection of age-related changes.

One view is that ageing leads to increased activity across the cortex due to dedifferentiation of cognitive mechanisms (Cabeza, 2002). Shih (2009) proposed that functional decline could be a result of impaired neural inhibition. Both decreased precision and inhibition could result in increased activation thresholds to select visual stimuli, thus, resulting in enhanced difficulties in reaching these thresholds (Adamo et al., 2003; Aydin et al., 2013; Huettel et al., 2001). The notion of inhibition has been strongly linked to alpha oscillations (8-12 Hz), including task-related modulations in amplitude and phase. (Capotosto et al., 2009; Hanslmayr et al., 2007; Hanslmayr et al., 2005; Klimesch et al., 2007; Rohenkohl and Nobre, 2011; Sauseng et al., 2005; Thut et al., 2006; Yamagishi et al., 2003). In addition to inhibition of irrelevant sensory information, alpha increases are also typically present during sustained (temporal) attention (Dockree et al., 2007; Rihs et al., 2007, 2009) and are likely to inhibit unattended locations and irrelevant sensory information (Bonnefond and Jensen, 2015; Busch et al., 2009; Dugué et al., 2011; Jensen and Mazaheri, 2010; Mathewson et al., 2009; Rihs et al., 2007). It has been reported that older adults do not modulate alpha oscillations to the same extent as younger adults (Deiber et al., 2013; Hong et al., 2015; Pagano et al., 2015; Vaden et al., 2012). This seems to be particularly the case in anticipation of a visual target (Deiber et al., 2013; Zanto et al., 2010), which could be indicative of a failure to inhibit irrelevant visual distractors (Vaden et al., 2012). However, reduced modulation of alpha oscillations does not seem to consistently result in impaired performance. Older individuals have been found to successfully inhibit visual information despite a lack of alpha modulation (Vaden et al., 2012), possibly indicating the implementation of alternative neural mechanisms, while alpha might become a mere indicator for progressing de-differentiation. However, the aforementioned research that presents alpha oscillations as a primary candidate for attentional gating has predominantly been conducted with young adults, mostly under the age of 30 years. It is therefore unclear to what extent attentional processes in young adults generalise to attention mechanisms in older participants. The present study set out to shed further light on whether changes in alpha oscillations could be indicative of de-differentiation or reduced inhibition in older age groups and explain deficits in attentional focusing.

In support of alternative processing strategies and more widely distributed brain activity in older age, there is cognitive evidence to suggest that older adults are able to compensate for attentional deficits with top-down control of attention, such as utilising cues more effectively than younger people in selective attention tasks (McLaughlin and Murtha, 2010; Neider and Kramer, 2011; Watson and Maylor, 2002). As proposed by the “Scaffolding Theory of Aging and Cognition” (STAC; Park and Reuter-Lorenz, 2009; Reuter-Lorenz and Park, 2014), successful compensatory cognitive strategies are likely to recruit additional neural resources, which could be reflected by a wider distribution of brain activity – prominently involving brain areas related to top-down control. Accordingly, the “posterior to anterior shift in ageing hypothesis” (PASA; Davis et al., 2008) proposes that there is a compensatory shift in activity towards frontal regions in conjunction with declines in occipital sensory processing, which has accrued supporting evidence (Buckner et al., 2000; Cabeza et al., 2004; Davis et al., 2008; Grady, 2000; Huettel et al., 2001; Madden, 2007; Madden et al., 2002; Ross et al., 1997). Crucially, increased frontal activity has been shown to correlate with decreased occipital activity (Cabeza et al., 2004; Davis et al., 2008) and improved task performance (Davis et al., 2008; Madden, 2007). A widely reported decline in the structure of frontal regions with age makes the PASA hypothesis somewhat counterintuitive. Colcombe et al. (2005), however, found that areas with the largest grey matter reductions, e.g. middle frontal gyrus (MFG) and superior frontal gyrus (SFG), also showed the greatest increases in activity. Similar patterns have been found in the ageing memory literature, where older people with hippocampal atrophy and episodic memory decline show the greatest increases in frontal lobe recruitment (Pudas et al., 2017).

However, inconsistent with a simple formulation of the PASA hypothesis of ageing (Davis et al., 2008), theta modulations (3-7hz) along the frontal midline have been reported to diminish with increasing age - in both resting-state and task-related conditions (Cummins and Finnigan, 2007; Reichert et al., 2016; van de Vijver et al., 2014). Theta oscillations are associated with a broad array of tasks measuring executive function and cognitive control (Cavanagh et al., 2009; Cavanagh and Frank, 2014; Demiralp and Başar, 1992; Green and McDonald, 2008; Min and Park, 2010; Sauseng et al., 2010). Frontal midline theta in particular is thought to reflect medial prefrontal cortex (mPFC) and, specifically, anterior cingulate cortex (ACC) activity (e.g. Asada et al., 1999) during attentional control (Cavanagh et al., 2009; Cavanagh and Frank, 2014; Konishi et al., 1999; Pollmann, 2004). Age-related reductions in frontal midline theta have most commonly been observed in memory recall tasks and during resting state, and were mostly recorded with electroencephalography (EEG; Cummins and Finnigan, 2007; Reichert et al., 2016; van de Vijver et al., 2014). There is evidence to suggest that theta power decreases from childhood throughout adulthood, which could reflect increased experience and reduced cognitive effort, however, with renewed increases later in life (Gómez et al., 2013). Consistent with these findings and a PASA hypothesis of ageing (Davis et al., 2008), Gazzaley et al. (2008) found an increase in frontal midline theta power in older adults when implementing a visual attention task, which could reflect an increase in the implementation of top-down attentional guidance. However, it remains unclear whether such increased activity is beneficial for performance or rather a further indication of de-differentiation and lack of neural precision.

In the light of the aforementioned inconsistencies and competing theoretical accounts, we set out to clarify whether impaired attentional control (refocusing from a temporal to a spatial task) in older adults (Callaghan et al., 2017) is characterised by an increased spread of activation or a reduced activation across cortical networks. Based on the reviewed findings, our primary focus of investigation was centred on modulations of alpha and theta frequency bands. We used Magnetoencephalography (MEG) to increase spatial resolution over previous EEG studies, while achieving the necessary temporal resolution for frequency-specific analysis, thus, allowing for oscillatory analysis in source space. The aim of the current study was to investigate the oscillatory patterns that reflect age-related changes in the ability to switch from allocating attention in time, to allocating attention spatially. Specifically, we compared age groups on their ability to identify a single target in a spatially focal but temporally changing RSVP stream, to identify a target in a spatially distributed but temporally unchanging VS display. We hypothesised that there could be an increase in frontal theta activity either reflecting beneficial, additional top-down processing (Davis et al., 2008; Fabiani et al., 2006; Gazzaley et al., 2008; Madden, 2007) or merely reflecting de-differentiation (Cabeza, 2002). The former would be reflected in an improvement in performance with increased theta modulation. Conversely, in the latter, no such improvement in performance would be observed with increased cortical recruitment. Alternatively, reduced midline theta power, as demonstrated in previous EEG studies (Cummins and Finnigan, 2007; van de Vijver et al., 2014), could be observed as a result of increased activation thresholds due to decreased neural precision (Huettel et al., 2001; Quandt et al., 2016; Welford, 1981). Based on the reviewed literature it was also expected that older adults would display abnormal alpha modulation, either through a weaker alpha power decrease (Deiber et al., 2013; Zanto et al., 2011), that could be indicative of reduced target processing, and/or through a weaker alpha power increase (Vaden et al., 2012), that could be indicative of reduced distractor suppression.

## Methods

### Participants

Participants were recruited from Aston University staff and students and the community. Participants aged over 60 years were also recruited from the Aston Research Centre for Healthy Ageing (ARCHA) participation panel. Participants provided written informed consent before participating and were screened for contraindications to having an MRI or MEG scan and received standard payment according to local rules. The research was approved by Aston University Research Ethics Committee (#776) and complied with the Declaration of Helsinki.

Sixty-three participants in three age groups (19-30, 40-49, 60+ years; see Table 1 for details) were included in the final analysis. Participants with visual impairments, photosensitive epilepsy, and a history of brain injury or stroke were excluded from participation. All participants in the 60+ years group (60-82 years) scored over the 87 cut-off for possible cognitive impairment on the Addenbrookes Cognitive Examination 3 (ACE-3; Noone, 2015). The ACE-3 consists of a series of short tasks that provide measures of language, memory, attention, fluency and visuospatial abilities. In total 73 participants were tested, but six participants were excluded from analysis due to low performance accuracy and/or too noisy MEG data resulting in fewer than 30 out of 80 trials remaining for one or more conditions after data pre-processing. These six participants included one individual aged 40-49 years and five participants aged 60+ years. Two participants withdrew from the study and in two data sets there was a recording error. Demographics for the remaining 63 participants are presented in Table 1.

**Table 1.** Participant demographics

|  |  | **Age Group (years)** |  |  |
| --- | --- | --- | --- | --- |
|  |  | 19-30 (n=20) | 40-49 (n=20) | 60+ (n=23) |
| **Age (years)** | Mean | 24.6 | 44.95 | 68.61 |
|  | SD | 2.96 | 3.28 | 5.43 |
| **Sex** | Male | 08 | 07 | 10 |
|  | Female | 12 | 13 | 13 |
| **Handedness** | Right | 16 | 19 | 22 |
|  | Left | 04 | 01 | 01 |
| **ACE-3** | Mean | n/a | n/a | 95.5 |
|  | SD | n/a | n/a | 2.69 |

This table presents the demographics for each age group, including participants' mean age, the number of participants who are male and female, the number of participants who are left and right handed, in addition to the mean ACE-3 scores for the 60+ years group.

### Materials and procedures

The attention switching paradigm from (Callaghan et al., 2017) was adapted for use with MEG (see Figure 1). The major change to the MEG paradigm was to reduce the number of conditions while increasing the number of trials in each condition (for the required signal-to-noise ratio for MEG analysis), by focusing only on pop-out VS, since Callaghan et al. (2017) had reported ceiling effects for Switch-Costs in serial VS. On each experimental trial participants attended to an RSVP stream first before switching to a pop-out VS display. Each trial consisted of a fixation cross, presented for 2000ms, followed by the RSVP stream, which was immediately followed by the VS display. E-Prime 2.0 Professional (Psychology Software Tool. Inc.) was used on a windows PC to present stimuli, record responses, and send triggers to the MEG through a parallel port (at the onsets of RSVP, target (if applicable), and VS display, as well as upon response to VS). Stimuli were back-projected onto a screen inside a magnetically shielded room (MSR) approximately 86cm in front of the participant at a resolution of 1400×1050. All stimuli were presented in black (RGB 0-0-0) on a grey background (RGB 192-192-192).

**Figure 1.**
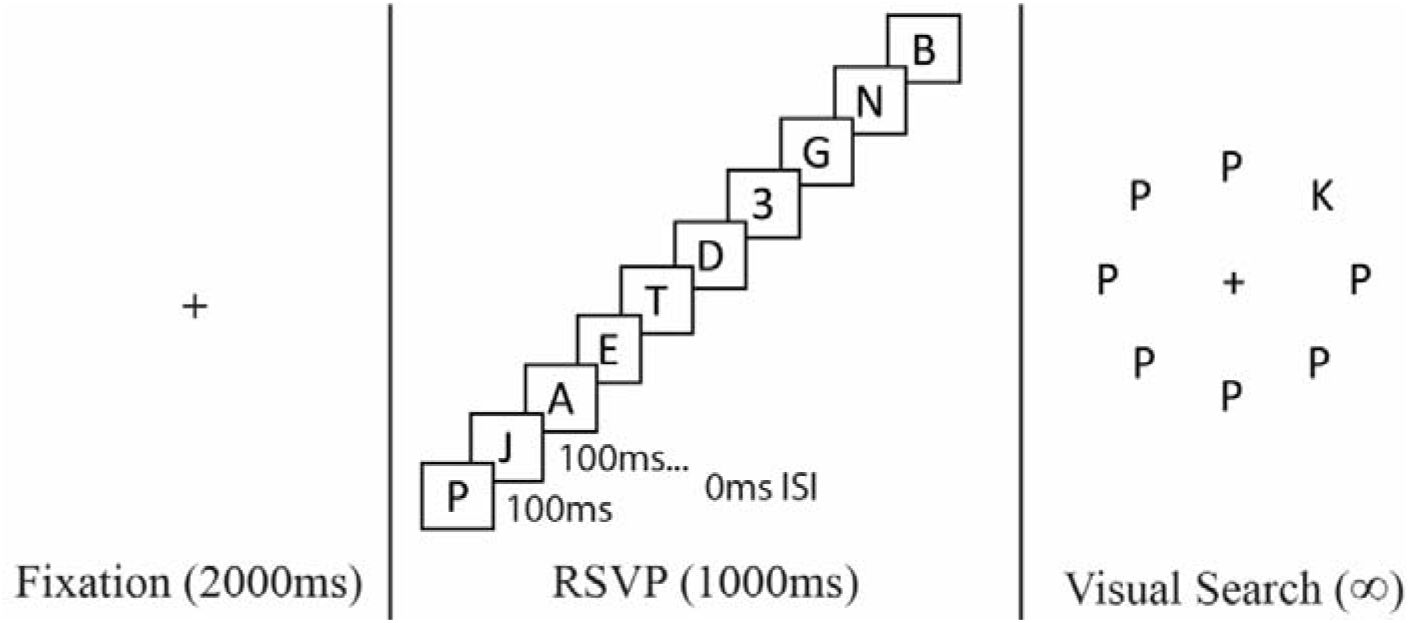
Illustration of trial structure and stimulus examples. The RSVP stream illustration (left) displays a Target Switch RSVP stream (a target digit at position 7 in the RSVP). Each trial consisted of a fixation cross (2000ms) followed by an RSVP stream immediately followed by a pop-out VS display (right). *ISI Inter stimulus interval*.

The RSVP stream consisted of a rapidly changing stream of letters in the centre of the display. There were ten items in each RSVP stream, each presented for 100ms with no inter-stimulus interval. Stimuli were presented in font size 30pt (0.75×0.75cm, 0.78°). On two thirds of the trials, one of the items in the stream was a target, namely a digit (1/2/3/4/6/7/8/9), which participants were expected to detect and memorise for report at the end of the trial (after the VS). The target could be either the first stimulus of the stream (removing the need to attend to the stream) or the seventh or ninth item in the stream of ten stimuli. In the remaining one third of the trials the RSVP contained only letters and no target digit. Due to its visual similarity to the letter S, ‘5’ was excluded from the pool of targets. Based on their visual similarity to certain numbers, letters I, O, and S were excluded from the stream. Letters K and Z were targets defined for the VS task and were therefore also not employed as distractors in the RSVP. It should be noted that the current RSVP task differs from a standard attentional blink paradigm as the RSVP stream only contained a maximum of a single target.

The VS display consisted of eight letters presented in a circle around a fixation cross in the centre of the screen, including seven distractors and one target. Participants were instructed to keep their eyes fixed on the cross at the centre of the screen while they completed the VS and to respond as quickly as possible. The target letter was always either a ‘K’ or a ‘Z’ and distractors were always a ‘P’, rendering a “pop-out” VS, conforming to effects observed by Callaghan et al. (2017; see Introduction for details). Stimuli were presented in font size 20pt (0.50×0.50cm, 0.52°) and the centre of each stimulus was 2.3cm (2.40°) from the centre of the fixation cross. Participants pressed a button with their right index finger once they had identified the VS target. Note that conforming to Callaghan et al. (2017) this button press did not discriminate between K or Z, but merely indicated that the participant had identified the target on that trial. Participants’ RTs to press this button were recorded and allowed for a more accurate and less variable search time estimate than a discriminative response (for detailed discussion, see Callaghan et al., 2017). For MEG it had the added benefit that this response did not trigger different neural motor patterns (e.g. for different finger taps). Subsequently, participants pressed a button to indicate whether it was a ‘K’ (right index finger response) or a ‘Z’ (left index finger response) in the display. Participants were then prompted to indicate whether they had seen a target digit in the RSVP stream (yes: right index finger response; no: left index finger response). If a digit was correctly detected in the RSVP stream, participants then pressed the button that corresponded with the number that they saw. Participants wore earphones through which a ‘ding’ sound was played after a correct response and a chord sound was played after an incorrect response.

To manipulate the cost of switching, the position of the target in the RSVP stream that preceded the VS was either the first item in the stream (Target No-Switch condition) or the target was either the seventh or ninth item in the stream (Target Switch condition) or absent from the stream (No-Target Switch condition). Illustrations of the RSVP stream and of the VS display are presented in Figure 1. There were 80 trials of each of the three conditions (No-Switch/Target Switch/No-Target Switch), with a total of 240 trials. To provide the opportunity for breaks, trials were divided into ten blocks. Trials were randomized within blocks. Participants completed 24 practice trials before starting the experimental trials.

MEG data were recorded with a 306-channel Elekta Neuromag system (Vectorview, Elekta, Finland) in a magnetically shielded room at a sampling rate of 1000Hz. The 306 sensors were made up of 102 triplets incorporating one magnetometer and two orthogonal planar gradiometers. Data were recorded in two halves within the same session.

Head position was recorded continuously throughout data acquisition via the location of five HPI coils. Three HPI coils were positioned across the participant’s forehead and one on each mastoid. The position of each HPI coil, three fiducial points, and 300-500 points evenly distributed across the head surface were recorded prior to the MEG recording with Polhemus Fastrak head digitisation. A T1 structural MRI was obtained for each participant, acquired using a 3T Siemens MAGNETOM Trio MRI scanner with a 32-channel head coil.

## Data analysis

### Response Times

Participants’ median VS RTs (ms) on trials where both VS and RSVP responses were correct were extracted. Participants’ proportions of correct VS target identifications and RSVP target identifications were also extracted.

Differences in median VS RTs between age groups and RSVP conditions were analysed in a 3 × 3 mixed ANOVA, where RSVP condition (No-Switch/Target Switch/No-Target Switch) was a within subjects factor and age group (19-30, 40-49, 60+ years) was a between subjects factor. Multiple comparisons were corrected for with Bonferroni correction.

The data were expected to violate assumptions of equality of variance due to increases in inter-individual variability with age (Hale et al., 1988; Morse, 1993), yet, there is evidence to support that ANOVA is robust to violations of homogeneity of variance (Budescu, 1982). Where Mauchly’s Test of Sphericity was significant, indicating that the assumption of sphericity had been violated, Greenhouse-Geisser corrected statistics were reported.

To interpret the age group × RSVP condition interactions, “Switch-Costs” were calculated as the percentage difference in RTs between Target Switch and No-Switch conditions (Target Switch-Costs) and between No-Target Switch and No-Switch conditions (No-Target Switch-Costs) for each individual. As interaction effects were already shown to be statistically significant in the ANOVA, Restricted Fisher’s Least Significant Difference test was applied and corrections for multiple comparisons were not conducted (Snedecor and Cochran, 1967). Where Levene’s test for equality in variance was significant (*p*<.05) when computing t-tests, ‘Equality of variance not assumed’ statistics were reported.

### MEG

MEG data were preprocessed in Elekta software using MaxFilter (temporal signal space separation, tSSS, .98 correlation; Taulu and Hari, 2009) to remove noise from sources inside and outside the sensor array. Seventeen participants displayed magnetic interference from dental work and so a tSSS correlation of .90 was applied instead. This included five participants from the 19-30 years group, six from the 40-49 years group and six from the 60+ years group. Movement correction was applied to one participant in the 40-49 years group due to head movement (>7mm).

Data were read into the Matlab® toolbox Fieldtrip version 20151004 (Oostenveld et al., 2011), with Matlab® 2015a, band-pass filtered between 0.5 - 85Hz and epoched from 3.5s preceding VS onset (i.e. 2.5s preceding RSVP stream onset) to 2.0s after the onset of the VS display. Trials were visually inspected for artefacts and any noisy trials were removed. Fieldtrip version 20161031 was used for further analysis.

Trials with incorrect responses were excluded. After excluding inaccurate and noisy trials, the mean number of trials that remained for each condition was 68.93 (SD=6.58) for the 19-30 years group, 67.97 (SD=7.51) for the 40-49 years group and 60.36 (SD=9.85) for the 60+ years group. Participants with fewer than 30 trials were excluded from the analysis (see Methods: Participants section).

### Sensor level analysis

For data cleaning prior to sensor level analysis, noisy sensors were interpolated with the average of neighbouring sensors. Independent components analyses (ICA) were implemented for each participant, across all conditions, and components with eye blink or heartbeat signatures were removed from the data.

Time-frequency analysis was carried out on signals from the planar gradient representation of 102 gradiometer pairs using a Hanning taper from 2-30Hz (for every 1Hz), with four cycles per time-window in stages of 50ms. For each participant trials were averaged within each condition (No-Switch/Target Switch/No-Target Switch).

To investigate the direction of task-related changes in oscillatory power in each age group, thereby improving interpretability of subsequent source level effects, we compared “active” task periods (3-5Hz: 550-1550ms relative to RSVP onset; 10-14Hz: 450-950ms and 1000-1500ms relative to RSVP onset; see *Source level analysis* section for details) to a baseline period (3-5Hz: −1500 - −500ms; 10-14Hz: −1000 - −500ms). Conditions were collapsed to obtain the average across all conditions. Two-tailed dependent t-tests were carried out to compare the active task periods with a baseline period separately for each age group.

Multiple comparisons were corrected for using non-parametric cluster permutations (Maris and Oostenveld, 2007), with 2000 permutations (cluster alpha = .05).

### Source level analysis

For source localisation using spatial filters (beamformers), noisy sensors were excluded rather than interpolated and ICA was not implemented to remove eye blinks and cardio artefacts. Due to size restrictions of the MEG data file, each data set was recorded in two halves within the same session and were therefore MaxFiltered separately prior to concatenating the data, which could lead to different components being removed in each half of data (see MaxFilter details in above MEG section). To reduce potential artefacts due to applying MaxFiltering to the two halves of data separately, a principle components analysis was implemented to reduce data dimensionality to components that accounted for 99% of the variance. The participant remained in the scanner and the door to the MSR remained shut between recording the two halves of data.

Using an in-house Matlab script and Elekta software MRI Lab, individual MRIs were aligned with the sensor array, by aligning the individual’s MRI with the fiducial positions and head shape that were recorded with Polhemus Fastrak head digitisation. Individual single-shell head-models (5mm voxels) were created from the coregistered MRIs. Head-models were normalised to MNI space (Montreal Neurological Institute template).

To identify the cortical generators of sensor level frequency modulations, we extracted time-frequency tiles from the time frequency representations (TFRs) in Figure 3, selecting 3-5Hz with a time window of 550-1550ms (relative to RSVP onset), and 10-14Hz with a time window of 450-950ms and 1000-1500ms (relative to RSVP onset) for theta and alpha frequencies, respectively. Note that this does not inflate type 1 error rates, as selection was not made by contrasting conditions or age groups, but rather on the overall pattern across all conditions and groups. The overlap between alpha and theta frequency ranges was minimised in order to capture distinct processing. For alpha it was possible to select two separate time windows (theta did not allow for such temporal resolution) to capture the two distinct phases of each trial (temporal and spatial tasks). Frequency-band specific Dynamic Imaging of Coherent Sources (DICS; Gross et al., 2001) beamformers (spatial filters; 2% lambda regularisation) were calculated based on cross-spectral densities obtained from the fast-fourier-transform (FFT) of signals from 204 gradiometers using a Hanning taper, spectral smoothing of +/-2Hz and 2.0s of data padding. No baseline correction was applied and conditions were directly compared instead. Note that, although group differences were also present in the beta frequency band (15-25Hz), our hypotheses focused on alpha and theta bands based on the previous literature (see Introduction). Surface plots in Figures 5, 6 and 8–11 were plotted in BrainNet Viewer 1.63 (Xia et al., 2013).

Two-tailed dependent t-tests were carried out to compare each of the Switch conditions (Target Switch/No-Target Switch) with the No-Switch condition separately for each age group. Multiple comparisons were corrected for with non-parametric cluster permutations (Maris and Oostenveld, 2007). Second level analysis was carried out by comparing Switch-Costs at the group level (Bögels et al., 2014; Wang et al., 2016). For each participant the No-Switch condition was subtracted from each of the Switch conditions separately. These differences were entered into two two-tailed independent cluster permutation t-tests (2000 permutations) to compare age groups (19-30 years vs 40-49 years/19-30 years vs 60+ years).

To explore the relationship between behavioural performance and power changes in theta and alpha frequencies in the two older groups (to better understand cognitive decline), differences in power (at peaks of the main significant clusters of the source analysis) between Target Switch conditions and the No-Switch condition in theta and alpha power were entered into Spearman’s correlation analysis with behavioural RT Target Switch-Costs. As no significant age group differences were found in No-Target Switch-Costs, we focus only on correlations between power change in the Target Switch condition (compared to No-Switch) and Target Switch-Costs in RT. To investigate possible correlations between behaviour and residual activity in regions shown to be involved in younger but not older groups, power change at the younger group’s cluster peaks were additionally entered into the older groups’ correlation analyses. Bonferroni correction was used to adjust the level of alpha to control for the number of correlations (the number of correlations are described below).

For the correlation of theta power change with Target Switch-Costs, one (parietal) ROI was chosen from the source level analysis (presented in Figure 5) from the 19-30 years group (reflecting residual activity in the older groups), in addition to two ROIs from each of the older and middle-aged groups. This resulted in six correlations for theta power in total, with three for each (older) age group (i.e. 60+ years: parietal, MFG and temporal lobe; 40-49 years: parietal, ACC and occipital lobe).

For the correlation of Target Switch-Costs and alpha power change during the RSVP window, one (parietal) ROI was chosen from the source level analysis of the 19-30 years group, and one ROI was chosen for each of the older and middle-aged groups (from the source level analysis presented in Figure 8). This resulted in four correlations, with two for each (older) age group (i.e. 60+ years: parietal and STG; 40-49 years: parietal and posterior parietal).

For the correlation of Target Switch-Costs with alpha power change during the VS window, one (IFG) ROI was chosen from the source level analysis from the 19-30 years group, and one ROI was chosen for each of the older and middle-aged groups (from the source level analysis presented in Figure 10). This resulted in four correlations, with two for each (older) age group (i.e. 60+ years: IFG and cerebellum; 40-49 years: IFG and cerebellum). Coordinates for the selected peaks can be found in Table SM1–3 in the supplementary material.

## Results

### Response Times: Switch-Costs

All groups correctly identified over 96% of VS targets in all three conditions. Thus, no further analysis was carried out on VS accuracy. All groups correctly identified over 73% of RSVP targets in both RSVP conditions. RSVP accuracy was unrelated to the aims and hypotheses of the current study and no further analysis was carried out on RSVP accuracy. The proportion of correct RSVP target identifications in the two Target conditions are presented in Figure SM1 in the SM. Group means of participants’ median VS RTs are presented in Figure 2.

**Figure 2.**
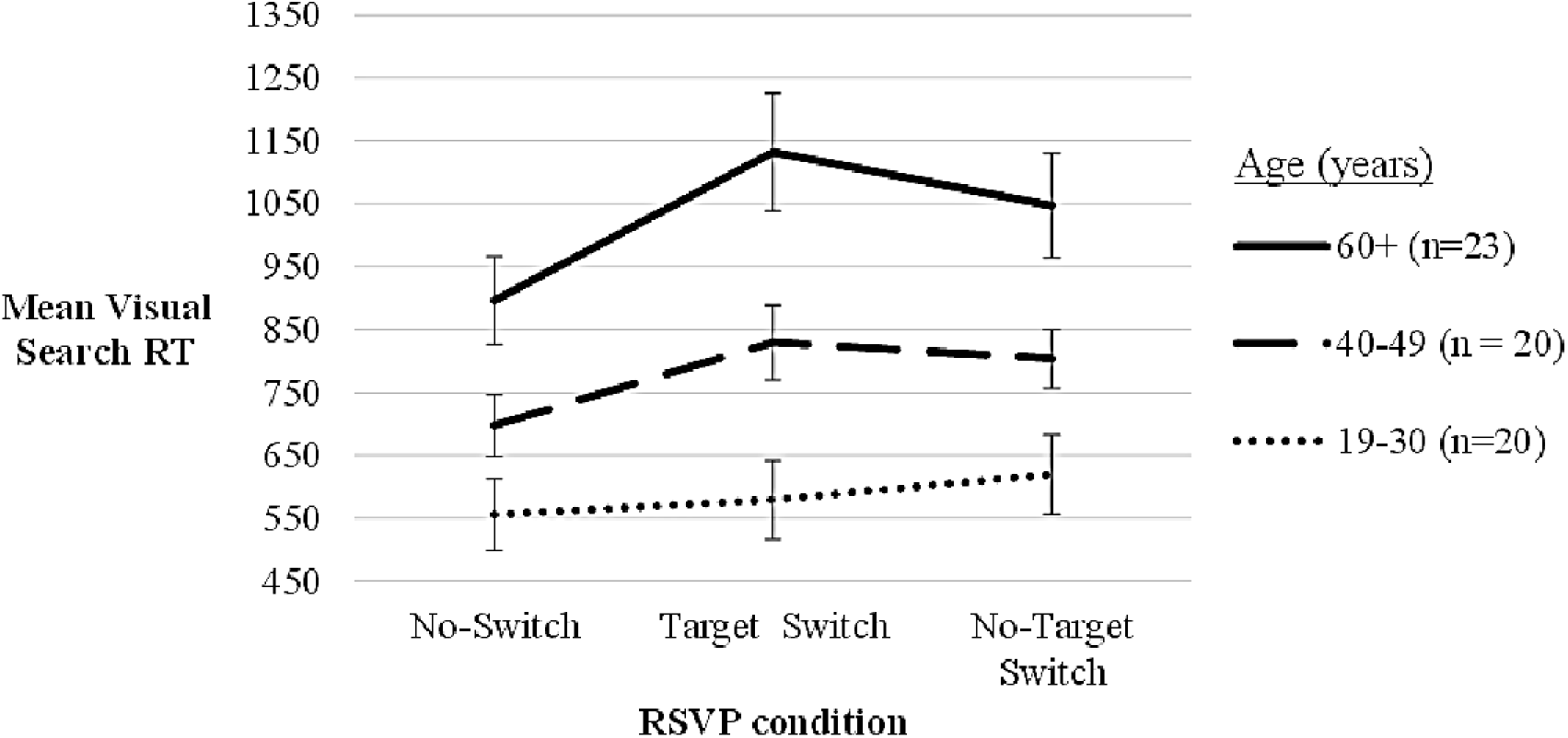
Group means of participants’ median VS RTs. Vertical bars represent the SE.

The 3 × 3 (RSVP condition × age group) mixed ANOVA on participants’ median VS RTs revealed a significant main effect of age (*F*(2, 60)=11.36, *p*<.001, η^2^_p_=.28), a significant main effect of RSVP condition (*F*(2,120)=35.21, *p*<.001, η^2^_p_=.37) and a significant interaction between age and RSVP condition (*F*(4,120)=7.05, *p*<.001, η^2^_p_=.19).

Post hoc comparisons revealed that the main effect of age resulted from significantly slower RTs in the 60+ years group in comparison to both the 19-30 (*p*<.001) and 40-49 years (*p*=.029) groups. There was no significant difference between the 19-30 and 40-49 years groups (*p*>.10).

The main effect of RSVP condition resulted from significantly slower RTs in both the Target Switch (*p*<.001) and No-Target Switch (*p*<.001) conditions in comparison to the No-Switch condition. There was no significant difference in RTs between the Target Switch and NoTarget Switch conditions (*p*>.10).

To investigate the hypothesis that there would be significantly greater Switch-Costs in both the 40-49 and 60+ years groups in comparison to the 19-30 years group, and to interpret the interaction between age and RSVP condition, independent t-tests were carried out comparing Switch-Costs across age groups. Please refer to Methods for a description of how Switch-Costs were calculated for each participant. Means and SDs of participants’ Switch-Costs are presented in Table 2.

**Table 2.** Means and SDs of Switch-Costs for each age group

|  |  | Age group (years) |  |  |
| --- | --- | --- | --- | --- |
|  |  | 19-30 | 40-49 | 60+ |
|  |  | (n=20) | (n=20) | (n=23) |
| **Target Switch-Costs** | Mean | 4.02 | 19.67 | 26.65 |
|  | SD | 12.72 | 15.78 | 15.67 |
| **No-Target Switch-Costs** | Mean | 12.59 | 17.29 | 17.98 |
|  | SD | 15.24 | 15.66 | 18.43 |

Target Switch-Costs were significantly greater in both the 40-49 ( $df=38$ , $t=-3.45$ , $p<.001$ ) and 60+ ( $df=41$ , $t=-5.15$ , $p<.001$ ) years groups in comparison to the 19-30 years group. There were no significant age group differences in No-Target Switch-Costs ( $p>.10$ ).

The RT results replicated findings from (Callaghan et al., 2017) by demonstrating deficits in switching in both the 40-49 years and 60+ years groups in comparison to the 19-30 years group. Consistent with Callaghan et al. (2017), greater Switch-Costs in the older age groups were only significant when participants were required to process a target digit before switching. When there was no target in the RSVP stream older participants seemed better able to switch from the temporal to the spatial attention task, suggesting either an increased demand for more processing resources and/or differences in strategies used to switch when target consolidation was required. To improve our understanding of the neurocognitive processing used to switch between modalities of attention across the three age groups, in the following sections we will investigate group differences in task related oscillatory signatures.

### MEG results

Frequencies from 2-30Hz were explored and TFRs are shown in Figure 3. Note that although group differences may also be present in the beta frequency band (15-25hz), our hypotheses focused on alpha and theta bands based on previous literature (see Introduction) and we therefore omitted beta in our analysis.

**Figure 3.**
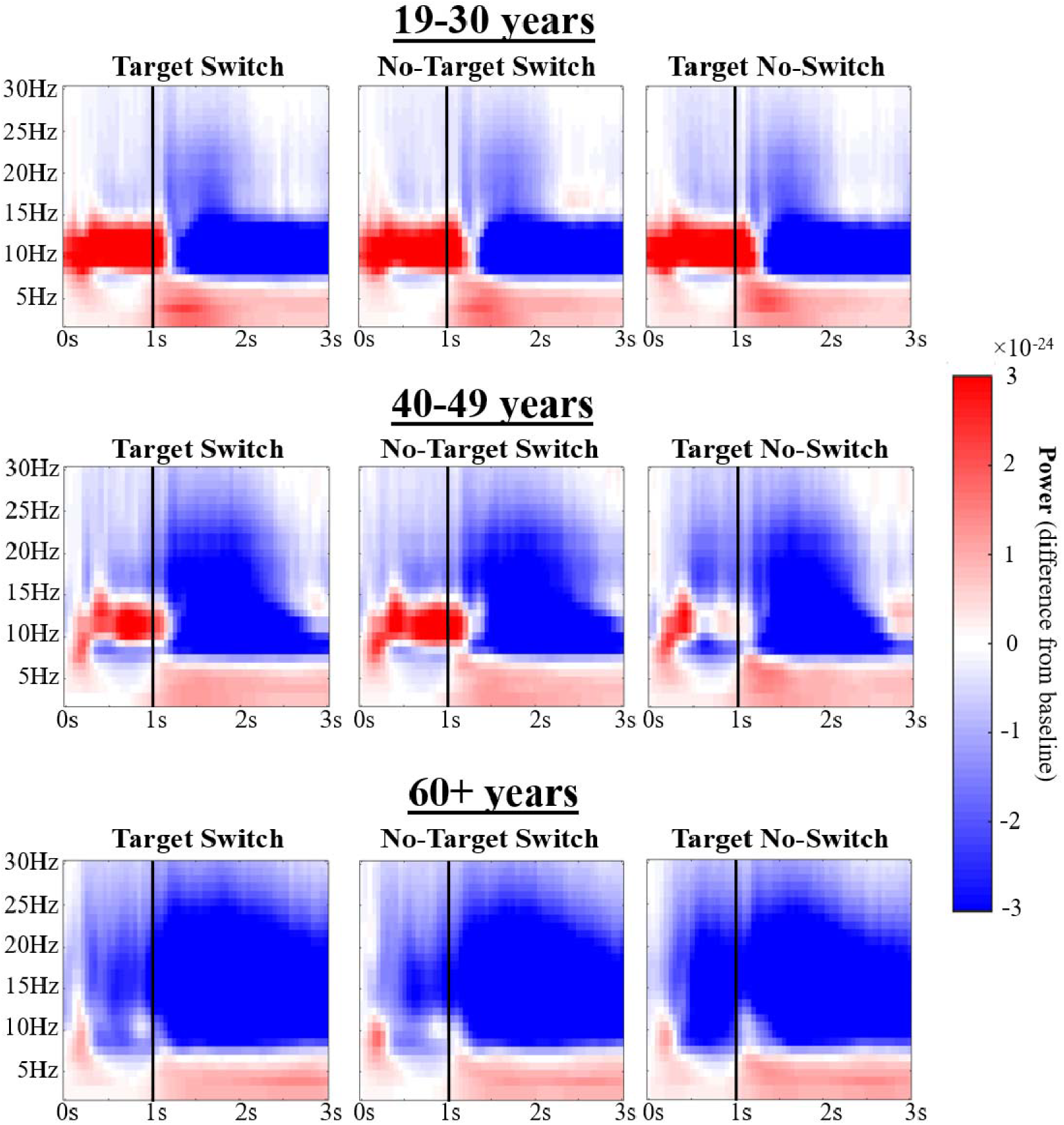
TFRs present power in relation to a baseline period of −0.6s - −0.01s in a group of four posterior gradiometer pairs. The onset of the RSVP stream occurred at 0.0s. Black lines placed over TFRs indicate the onset of the VS display, and RSVP target onset occurred at either 0.7 or 0.9s.

### Theta power in source space

**Figure 4.**
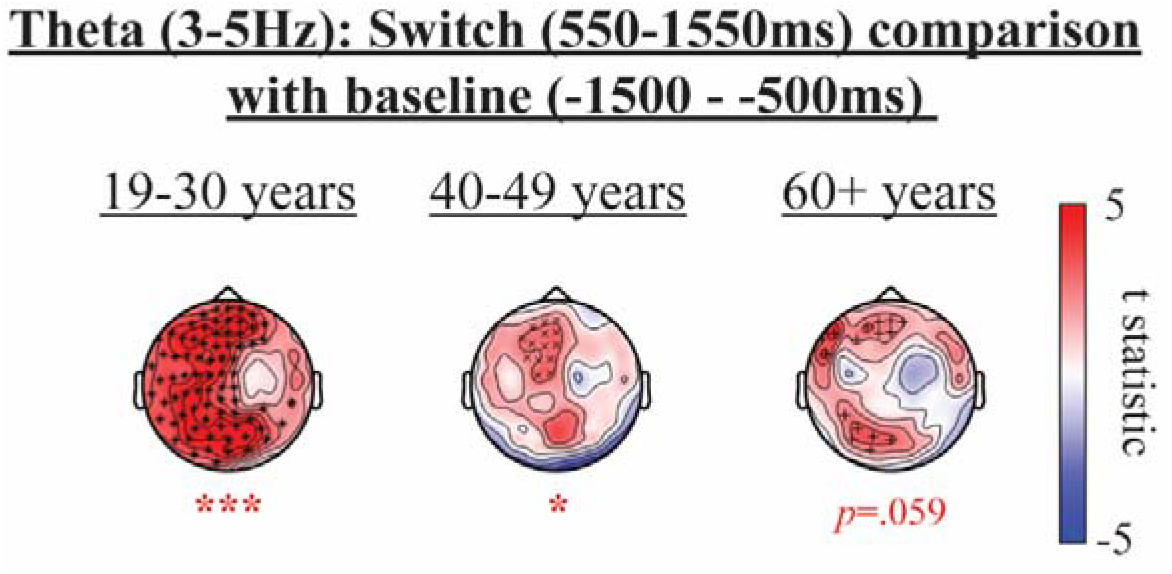
Effects in theta (3-5Hz) when contrasting Switch period (550-1550ms; collapsed across all 3 RSVP conditions) to the baseline period (−1500 - −500ms), for each age group. Sensor topographies present *t*-statistics of significant clusters (*p<.025, ***p<.001; positive clusters denoted in red).

Figure 4 shows a significant increase in theta power in relation to baseline, in a time window of 550-1550ms, in all age groups. Statistical results comparing theta power in Target Switch and Target No-Switch conditions and exploring the interaction between RSVP condition and age group, are presented in Figure 5 (for details see Methods, Section 2).

**Figure 5.**
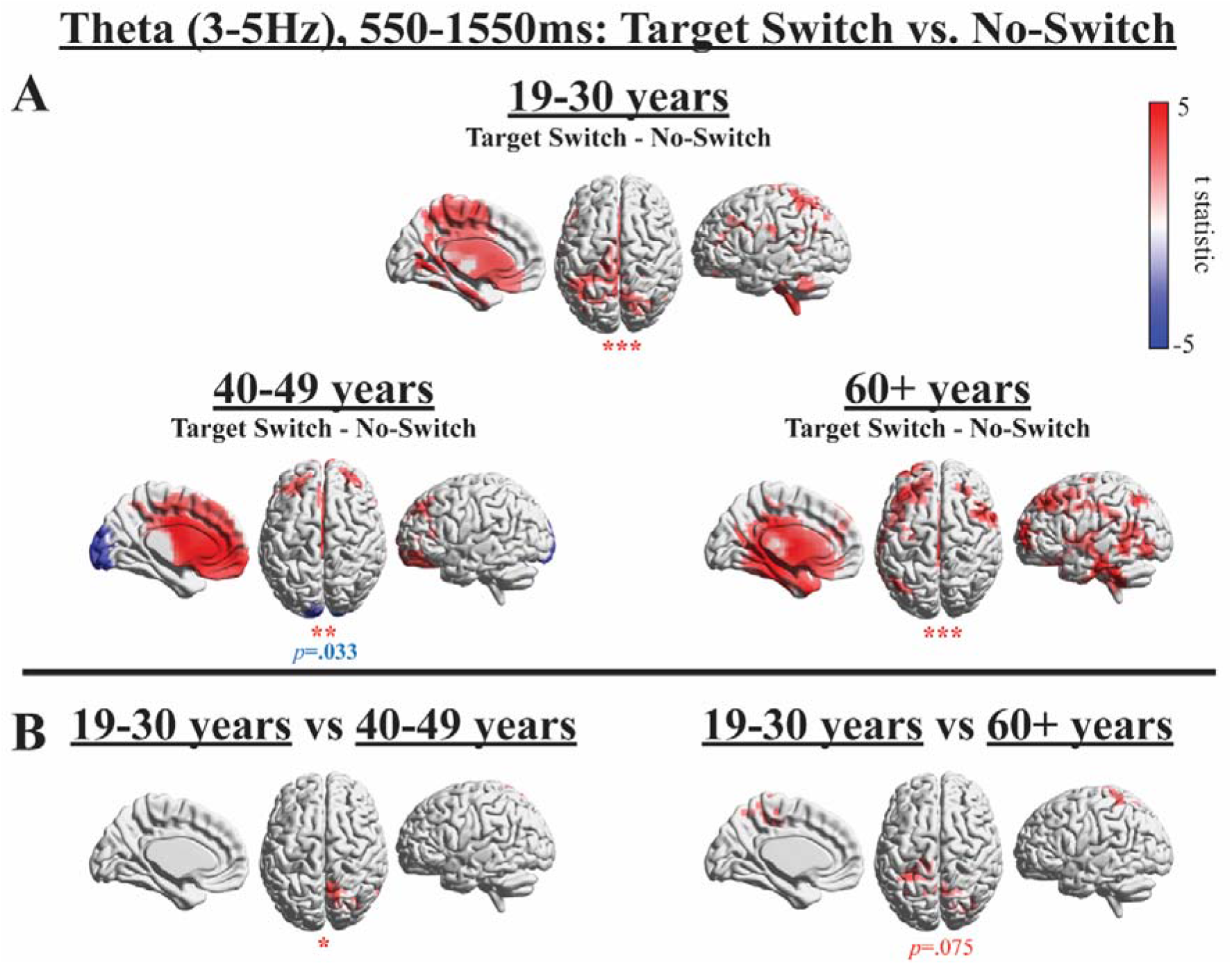
Effects in theta (3-5Hz) when contrasting Target Switch and No-Switch conditions in each age group (panel A) and when exploring the Target Switch condition × age interaction (panel B). The colour bar displayed in panel A applies to both panel A and B. Source plots present *t*-statistics of significant clusters (*p<.025, **p<.01, ***p<.001; positive clusters denoted in red; negative clusters denoted in blue). For unthresholded effects see Figure SM2 (Supplementary Material).

The TFRs in Figure 3 and sensor level analysis in Figure 4 illustrate that there was a theta increase in response to the VS display onset in all conditions. Figure 5 illustrates that all age groups displayed a significantly higher theta increase in the Target Switch condition in comparison to the No-Switch condition, which localised to superior and inferior parietal gyri, occipital gyri, and the MFG in the 19-30 years group, bilateral frontal cortex and the ACC in the 40-49 years group and the SFG, temporal gyri and the cerebellum in the 60+ years group (Figure 5, panel A). Whereas the 19-30 years group displayed higher theta in parietal regions, the two older groups demonstrated more extensive frontal recruitment. This increase in left MFG theta power was correlated with decreased Switch-Costs in the 60+ years group (*r*=-.40, *p*=.057). The 60+ years group additionally displayed higher temporal lobe theta that was not present in the two younger groups. The 40-49 years group additionally presented with a posterior (occipital/cerebellar) negative cluster, which reflects lower theta in the Target Switch condition in comparison to the No-Switch condition, although this did not reach significance in a two-sided test (*p*=.033). Note that the spread of source power to the centre of the brain in medial slices in Figures 5, 8 and 9 are a result of spatial leakage, a known challenge with MEG source analysis.

Age-group comparisons of differences between Target Switch and No-Switch conditions, which are presented in Figure 5 panel B, confirmed that the higher theta increase in the Target Switch condition was greater in the 19-30 years group in parietal regions in comparison to the 40-49 years group (*p*=.020) and the 60+ years group (*p*=.075), although the latter did not reach significance. In the 60+ years group, greater theta power increases in this parietal region were associated with decreased Target RT-Switch-Costs (r=-.53, *p*=.010). Importantly, due to reduced parietal theta in the 60+ years group overall (Figure 5, panel B), the coordinates for the parietal correlation effect were adopted from the 19-30 years group, in order to specifically investigate whether residual theta power in the oldest participants would be beneficial for attention switching. However, correlations did not survive a more stringent alpha level of *p*<.008 after Bonferroni correction to control for the number of tests performed (n=6). No such correlations were observed for the middle-aged group (*p*>.10 uncorrected).

**Figure 6.**
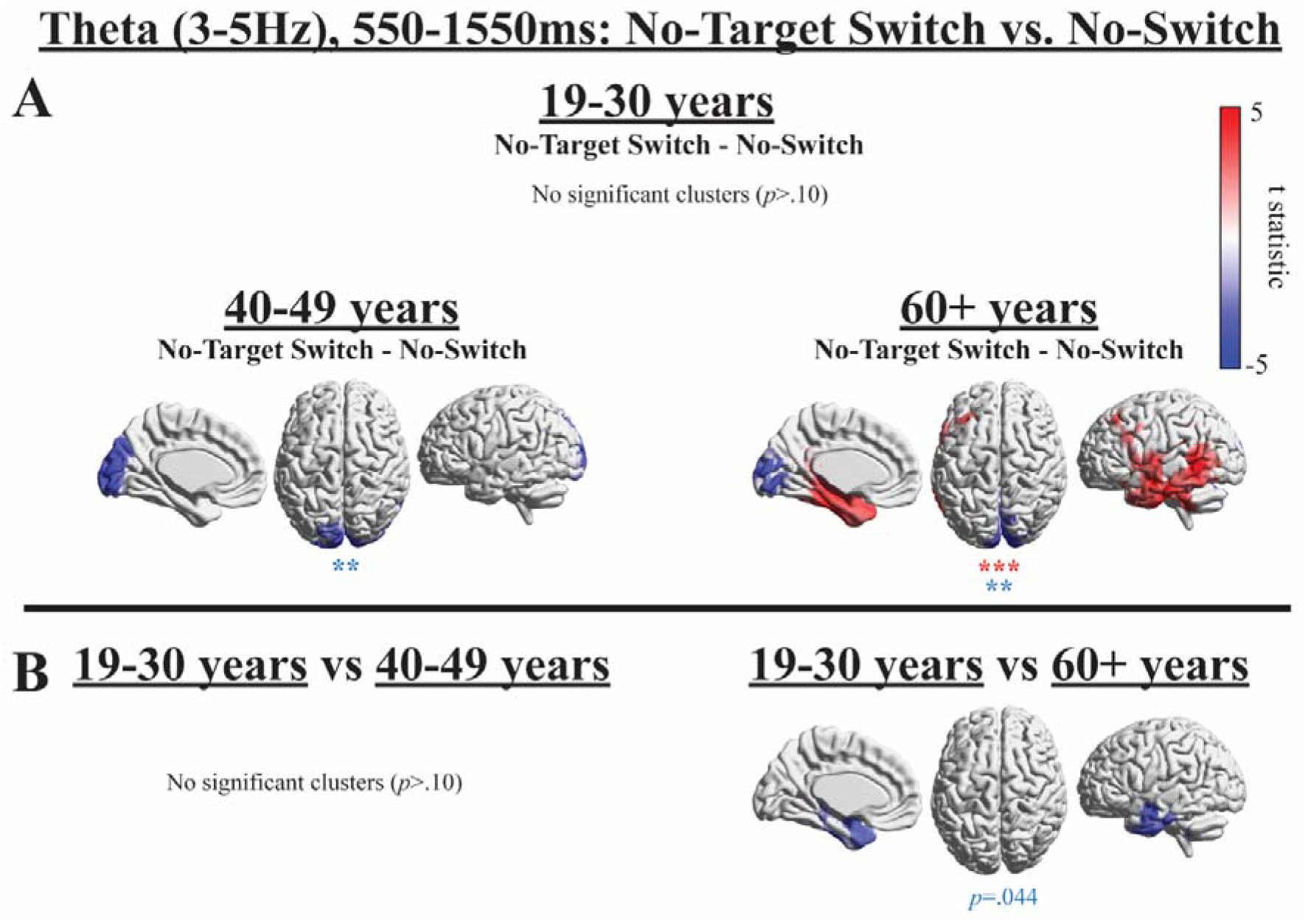
Effects in theta (3-5Hz) when contrasting No-Target Switch and No-Switch conditions in each age group (panel A) and when exploring the No-Target Switch condition × age interaction (panel B). The colour bar displayed in panel A applies to both panel A and B. Source plots present *t*-statistics of significant clusters (**p<.01, **p<.001; positive clusters denoted in red; negative clusters denoted in blue).

Figure 6A reveals that there was no significant difference between No-Target Switch and No-Switch conditions at theta frequency in the 19-30 years group, suggesting that the differences observed in theta between Target Switch and No-Switch conditions in this age group (see Figure 5) were a result of processing the RSVP target in the Target Switch condition.

In contrast, both the 40-49 and 60+ years groups again display negative clusters that localise to the occipital lobes, indicating deficient theta increases in the No-Target Switch condition, a finding that cannot be due to RSVP target processing. The 60+ years group again showed higher theta in the No-Target Switch condition in comparison to the No-Switch condition that localised to frontal regions and the left temporal lobe. However, group differences did not reach significance for a two-sided test (Figure 6, Panel B).

### Summary theta

In summary, the 19-30 years group showed higher theta power related to a Target Switch in parietal regions, an effect that was stronger compared to the two older groups (although, this difference did not reach significance in the 60+ years group). However, increased theta seems to be related to RSVP target processing, as no significant difference in theta was observed between No-Target Switch and No-Switch conditions in the 19-30 years group.

Contrary to the 19-30 years group, both the 40-49 and 60+ years groups showed significantly lower occipital theta in Target Switch (40-49 years) and No-Target Switch (40-49 & 60+ years) conditions (compared to the No-Switch condition). It could be that occipital theta deficits in the two Switch conditions are a reflection of deficient attentional guidance in visual processing regions, possibly contributing to the increased VS RTs observed in the two older groups after switching.

The 60+ years group additionally showed significantly higher frontal and temporal theta in the two Switch conditions in comparison to the No-Switch condition, and the 40-49 years group showed higher frontal theta in the Target Switch condition. It could be that this additional recruitment of the frontal cortex reflects the two older groups recruiting additional resources and relying more on top-down attentional control (McLaughlin and Murtha, 2010; Neider and Kramer, 2011; Watson and Maylor, 2002). The additional recruitment of temporal gyri in the 60+ years group may indicate the implementation of further strategies to cope with task demands, such as enhanced episodic memory encoding (Schacter and Wagner, 1999) or silent vocalisation (Graves et al., 2007; Hickok and Poeppel, 2007; Hocking and Price, 2009; Smith et al., 1998).

### Alpha power in source space

**Figure 7.**
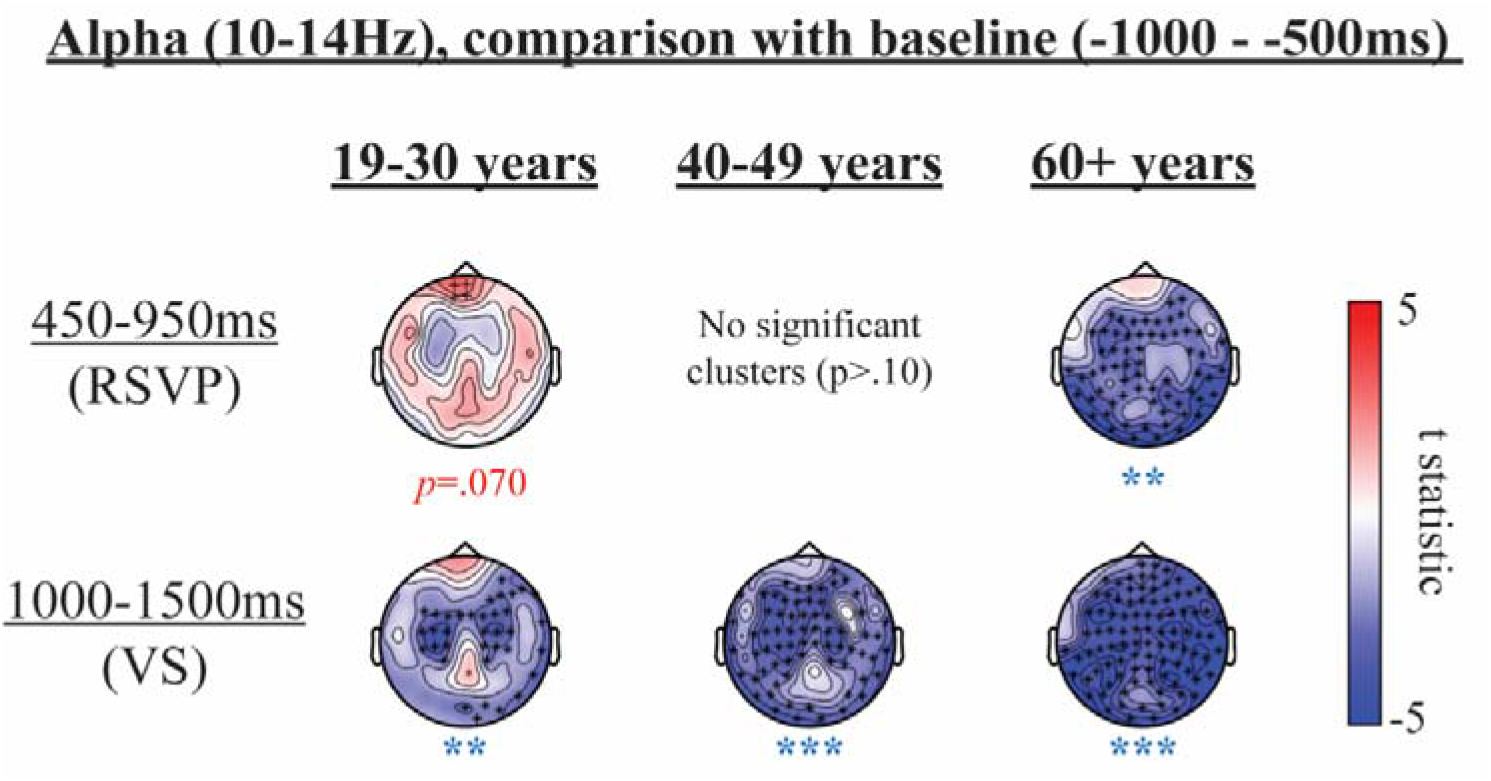
Effects in alpha (10-14Hz) when contrasting an RSVP window (450-950ms) and the VS onset window (1000-1500ms) to the baseline period (−1000 - −500ms), for each age group, collapsed across conditions. Sensor topographies present *t*-statistics of significant clusters (**p<.01, ***p<.001; positive clusters denoted in red; negative clusters denoted in blue).

There was a non-significant increase in alpha power in relation to baseline in the 450-950ms time window (relative to RSVP onset) in the 19-30 years group (see Figure 7; also Figure 3). In contrast, the 60+ years group showed a significant decrease in alpha power in relation to baseline in the same time window, and the 40-49 years group showed no significant difference (see Figure 7; also Figure 3). There was a significant decrease in alpha power compared to baseline in the VS time window (1000-1500ms relative to RSVP onset) in all age groups (see Figure 7; also Figure 3). Figure 8 presents the statistical results that compare alpha power in Target Switch and No-Switch conditions (panel A), as well as the interaction between RSVP condition and age group (panel B).

**Figure 8.**
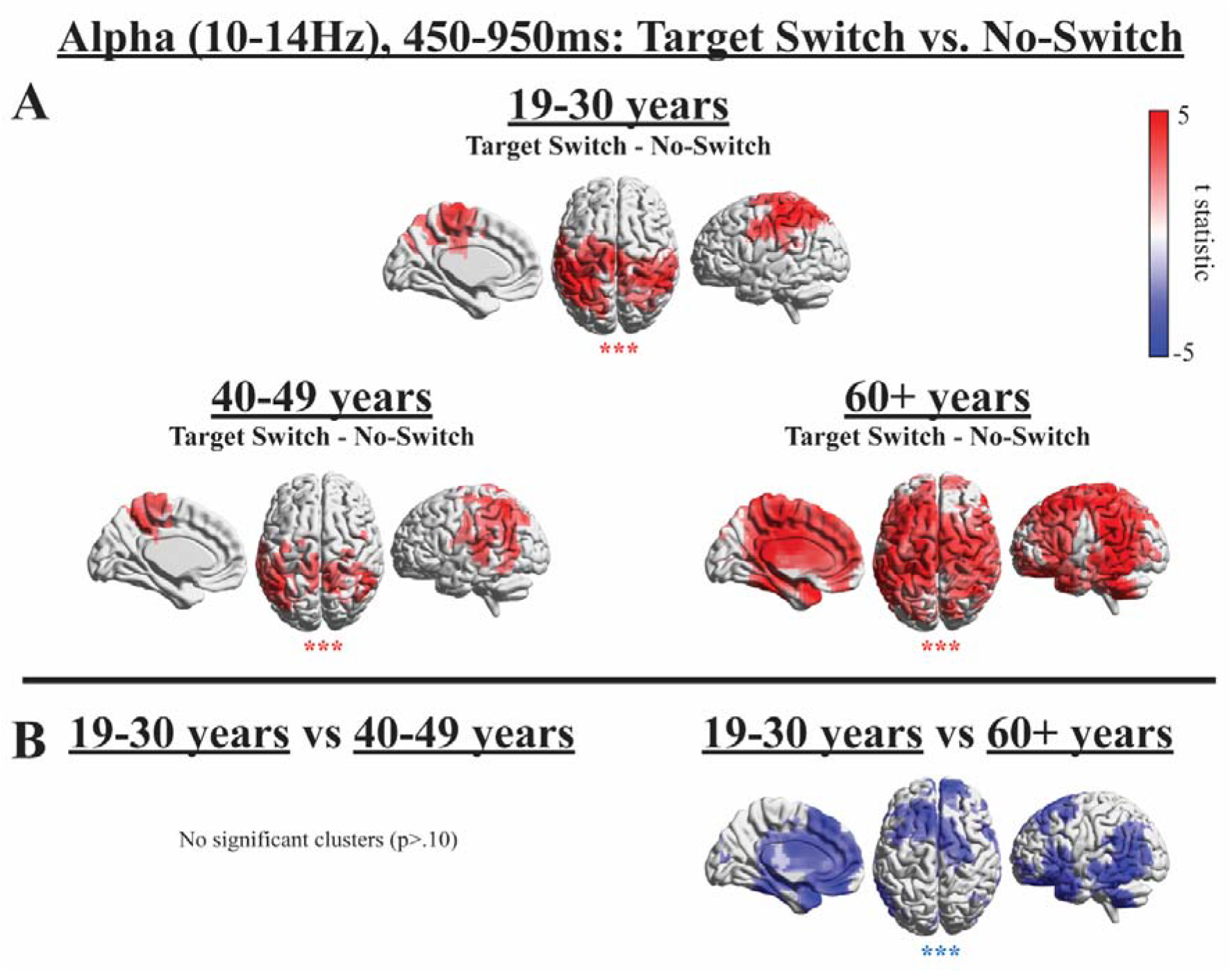
Effects in alpha (10-14Hz) during RSVP when contrasting Target Switch and No-Switch conditions in each age group (panel A) and when exploring the Target Switch condition × age interaction (panel B). The colour bar displayed in panel A applies to both panel A and B. Source plots present *t*-statistics of significant clusters (***p<.001; positive clusters denoted in red; negative clusters denoted in blue).

All age groups show significantly higher alpha power in the Target Switch condition in comparison to the No-Switch condition during the RSVP stream, which localised primarily to parietal regions in all age groups but was widely distributed across the cortex in the 60+ years group (Figure 8, panel A). In contrast to the 19-30 years group, both the 40-49 and 60+ years groups displayed higher temporal lobe alpha in the Target Switch condition in comparison to the No-Switch condition (Figure 8, panel A).

Figures 3 and 7 suggest that, in the 19-30 years group, this difference in alpha resulted from an alpha increase throughout the RSVP stream that was higher in the Target Switch condition than the No-Switch condition. In contrast, in the 60+ years group, higher alpha in the Target Switch condition resulted from a greater alpha decrease in the No-Switch condition than the Target Switch condition throughout RSVP presentation (Figure 3). Although no significant change in alpha power in relation to baseline was detectable in the 40-49 years group (see Figure 7), the TFRs displayed in Figure 3 suggest that this group’s pattern of alpha oscillations was closer to the younger group, suggestive of a greater alpha increase throughout the RSVP stream that was higher in the Target Switch condition than the No-Switch condition.

Group comparisons of differences highlighted that the higher alpha in the Target Switch condition in comparison to the No-Switch condition was significantly greater in the 60+ years groups in comparison to the 19-30 years group, as is reflected by the widely distributed negative cluster in Figure 8 panel B, spanning frontal, parietal, and temporal areas. However, the different origins of this group effect should be kept in mind, when interpreting the result, since younger participants revealed an alpha increase during the RSVP, while older participants presented with an alpha decrease (see Figures 3 and 7). There was no significant difference between the 19-30 and 40-49 years groups (*p*>>.10). There were no significant correlations between the change in alpha power at cluster peaks (during the RSVP window) and Target Switch-Costs (all *p*>.10 uncorrected). As stated above, the spread of source power to the centre of the brain in medial slices in Figures 5, 8 and 9 are a result of spatial leakage.

**Figure 9.**
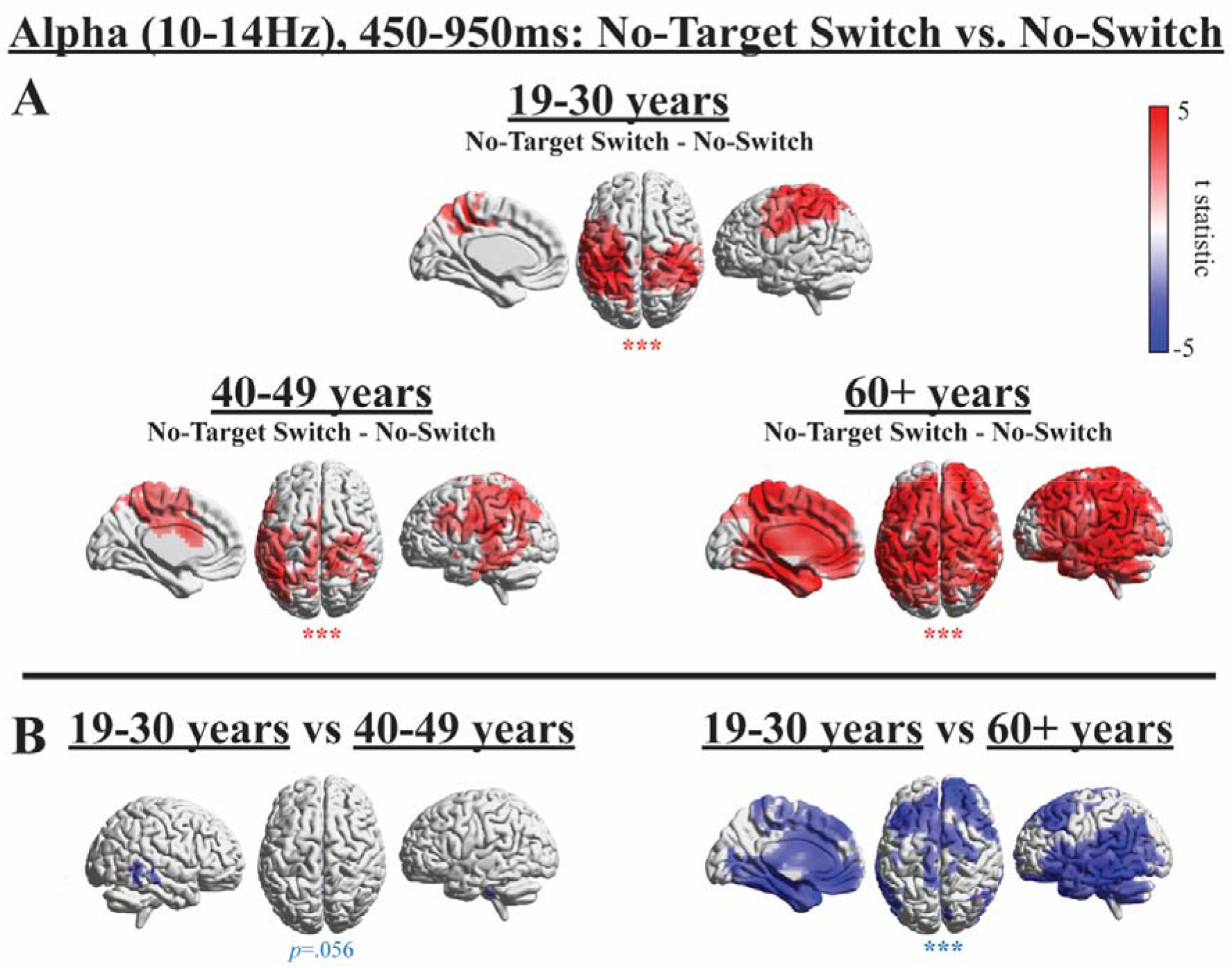
Effects in alpha (10-14Hz) during RSVP when contrasting No-Target Switch and No-Switch conditions in each age group (panel A) and when exploring the No-Target Switch condition × age interaction (panel B). The colour bar displayed in panel A applies to both panel A and B. Source plots present *t*-statistics of significant clusters (***p<.001; positive clusters denoted in red; negative clusters denoted in blue).

Similar to the Target Switch vs. No-Switch contrast, all age groups show significantly higher alpha in the No-Target Switch condition in comparison to the No-Switch condition in the RSVP time window, which localised to parietal regions in all age groups but was more widely distributed across the cortex in the 60+ years group. Similar to the pattern seen when comparing Target Switch and No-Switch conditions in Figure 8, lower alpha in the No-Switch condition in comparison to the No-Target Switch condition appears to have resulted from a greater alpha increase in the Target Switch condition in the 19-30 and 40-49 years groups and a greater alpha decrease in the No-Switch condition in the 60+ years group (see Figures 3 and 7), which is important to consider when interpreting intra- and inter-group effects.

Group comparisons revealed that the higher alpha in the No-Target Switch condition in comparison to the No-Switch condition was significantly higher in the 60+ years groups in comparison to the 19-30 years group, as is reflected by the negative clusters in Figure 9 panel B. Group differences between the 19-30 and 40-49 years groups did not reach significance (*p*=.056). While alpha effects were contained to parietal regions in the 19-30 years group, in the 40-49 and especially in the 60+ years groups the higher alpha effects were both stronger and more widely distributed across the cortex. In the 40-49 years group the distribution extended primarily into the ventral processing stream in occipito-temporal cortex, whereas in the 60+ years group the wider distribution also comprised frontal and prefrontal areas.

In response to VS onset, the 19-30 years group displayed a greater alpha decrease in the No-Switch compared to the Target Switch condition in frontal cortex (Figure 10 panel A). In contrast, the 40-49 years and 60+ years groups show a greater alpha decrease in the Target Switch condition compared to the No-Switch condition in parietal cortex and cerebellum. Group comparisons demonstrated that such group differences were significant (Figure 10 panel B). There were no significant correlations between the change in alpha power at cluster peaks (during the VS window) and Target Switch-Costs (*p*>.10 uncorrected).

**Figure 10.**
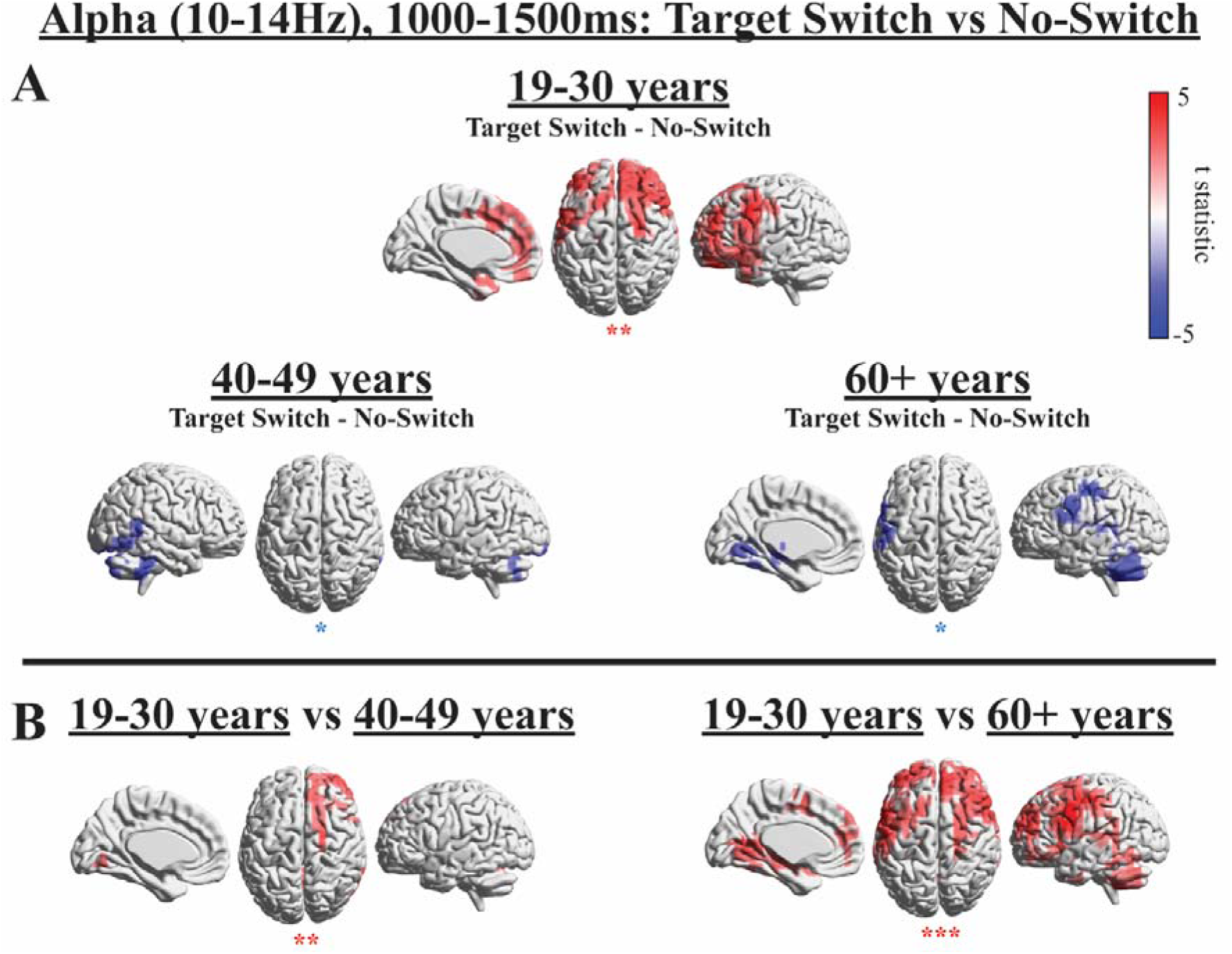
Effects in alpha (10-14Hz) during VS when contrasting Target Switch and No-Switch conditions in each age group (panel A) and when exploring the Target Switch condition × age interaction (panel B). The colour bar displayed in panel A applies to both panel A and B. Source plots present *t*-statistics of significant clusters (*p<.025, **p<.01, ***p<.001; positive clusters denoted in red; negative clusters denoted in blue).

In response to VS onset, both the 19-30 years and 60 + years groups displayed a greater alpha decrease in left frontal and parietal cortex in the No-Switch compared to the Target Switch condition (Figure 11 panel A). In contrast, the 40-49 years group show a greater alpha decrease in occipital cortex in the Target Switch condition compared to the No-Switch condition. The cluster in the 40-49 years group also extends to the cerebellum, however, the estimation of sources close to the edge of the sensor array can be poor and should be interpreted with caution, as this could be a result of spatial leakage. The group differences between the 19-30 years and 40-49 years groups did not reach significance in the group comparisons (Figure 11 panel B), nor did the greater alpha decrease in left frontal cortex in the No-Switch compared to the Target Switch condition in the 19-30 years group compared to the 60+ years group.

**Figure 11.**
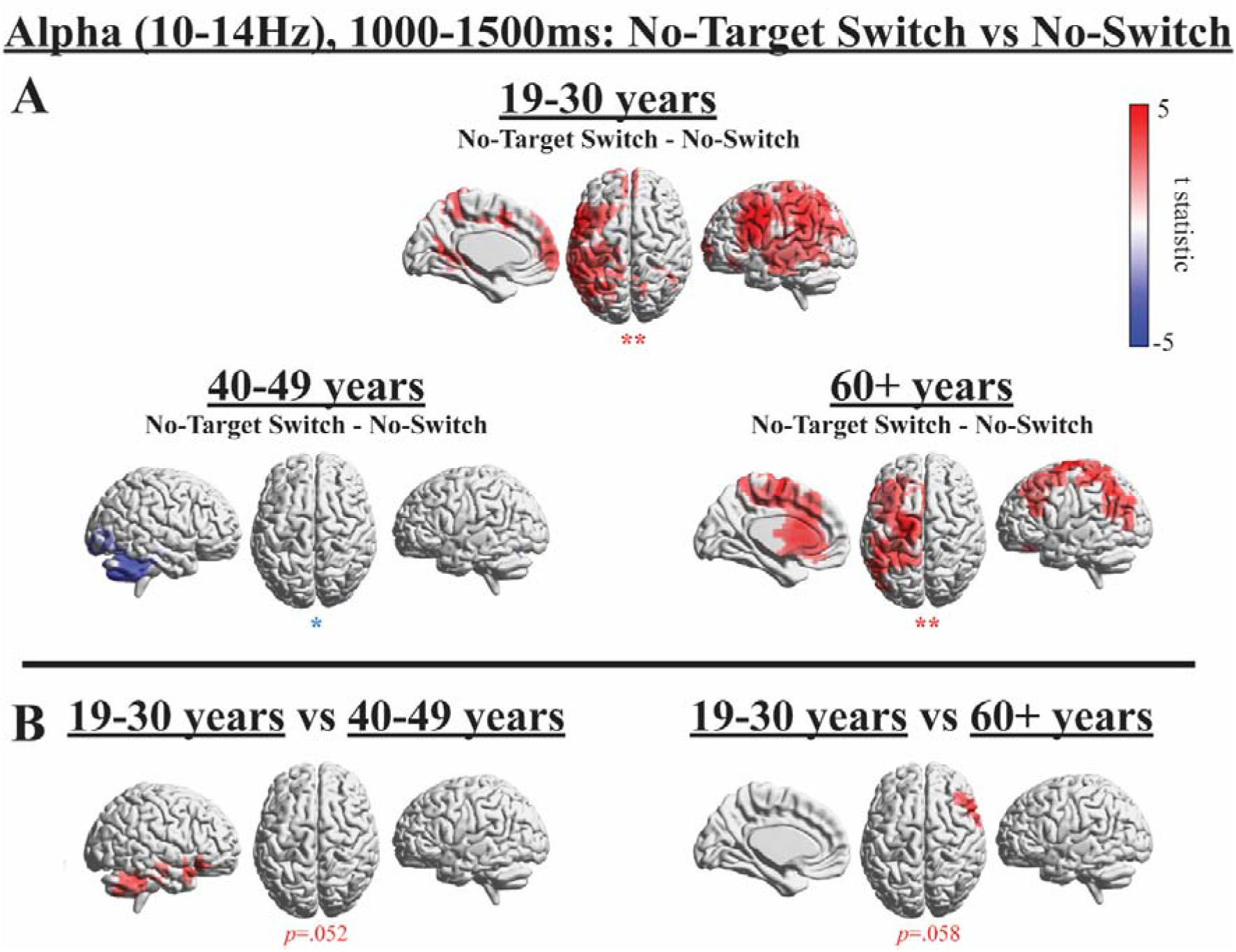
Effects in alpha (10-14Hz) during VS when contrasting No-Target Switch and No-Switch conditions in each age group (panel A) and when exploring the No-Target Switch condition × age interaction (panel B). The colour bar displayed in panel A applies to both panel A and B. Source plots present *t*-statistics of significant clusters (*p<.025, **p<.01; positive clusters denoted in red; negative clusters denoted in blue).

## Discussion

In our previous work we demonstrated that older adults find refocusing attention from time to space more difficult than younger adults (Callaghan et al., 2017). In the current study we replicated these results and found that the older (60+) as well as the middle-aged (40-49) group had increased Switch-Costs compared to the younger (19-30) group, as reflected by disproportionately increased RTs when required to refocus attention from a temporal RSVP task (when it included a target) to a spatial VS task. The primary aim of the current study was to investigate the age-related changes in neural mechanisms that may underlie this difficulty in refocusing attention from events changing in time to stimuli distributed spatially. We aimed to determine whether changes in attention refocusing are characterised by a reduced activation across cortical networks or an increased spread of activation, which could reflect either increased compensation or de-differentiation.

Also consistent with Callaghan et al. (2017), RTs of the 60+ years group were slower overall in comparison to the 19-30 years group. On the other hand, RTs of the 40-49 and 19-30 years groups did not significantly differ, implying that the 40-49 years group found the baseline No-Switch condition no more demanding than younger adults. However, the 40-49 years group again presented significantly higher Switch-Costs than the 19-30 years group, suggesting that they found the Target Switch condition disproportionality more demanding than the No-Switch condition, contrasting with the 19-30 years group. The 40-49 years group indeed seems to represent an intermediate stage of ageing, where some aspects of attentional control function at a similar level to younger adults, whereas other aspects coincide more with patterns observed in older adults, as observed in both RTs and neural oscillations.

Conforming to our hypotheses based on previous reports (Cummins and Finnigan, 2007; Deiber et al., 2013; Gazzaley et al., 2008; Vaden et al., 2012; van de Vijver et al., 2014), we indeed observed modulations of theta and alpha oscillatory power (Figures 3–11). The hypothesis that there would be reduced theta power with increased age was partially supported. The enhanced spatial resolution of MEG compared to EEG allowed us to go beyond the previous literature to investigate group differences in source space. We observed reduced theta power in occipital and parietal regions. However, instead of a reduction in frontal midline theta power, as indicated by several previous reports (Cummins and Finnigan, 2007; Reichert et al., 2016; van de Vijver et al., 2014), frontal midline theta was increased for the attention Switch conditions in the older and middle-aged groups in relation to a Target Switch. A relative increase in frontal midline theta (in relation to the No-Switch condition) with increased age is in line with the findings of Gazzaley et al. (2008). The 60+ years group presented with a more widely distributed relative theta increase in frontal regions across both Switch conditions. Furthermore, the observed correlation between reduced Switch-Costs and both frontal (MFG) and parietal theta (which was not significant after correction but had a medium effect size; Cohen, 1992) indicates that increased and more widely spread theta power is more likely to reflect alternative, yet beneficial processing, instead of mere de-differentiation (Cabeza, 2002) or lack of neural precision (Shih, 2009; Welford, 1981). Theta power findings therefore further support that older adults might recruit additional executive resources (Davis et al., 2008; Fabiani et al., 2006; Madden, 2007), as well as a PASA hypothesis of ageing (Davis et al., 2008), which proposes a posterior to anterior shift with increasing age.

As anticipated, there were age-related changes in task related alpha modulation during both the RSVP and VS time windows. During the RSVP window, instead of showing an alpha increase to inhibit irrelevant visual information (Vaden et al., 2012), the 60+ years age group showed an alpha decrease. This stronger and widely distributed alpha desynchronization (Figures 3 & 7–9) could reflect an enhanced attention strategy rather than an inhibition strategy. A lack of alpha synchronisation is in line with the hypothesis of reduced inhibition in older adults (Adamo et al., 2003). The middle-aged group presented an intermediate pattern at sensor level (Figure 3), where no significant difference in alpha power from baseline was detectable (collapsed across all conditions; see Figure 7), which differed from a significant alpha increase in the younger adults and a significant alpha decrease in the older adults. TFRs (Figure 3) seem to imply that the 40-49 years group was closer to displaying an alpha increase similar to younger adults, rather than an alpha decrease similar to older adults (Figure 3). Yet in source space, the pattern of switch-costs in the middle-aged group appeared to be more similar to the older group, with a stronger and more widely distributed alpha modulation across the cortex than the younger group. However, such differences did not reach significance in direct group comparisons (Figures 8 and 9), further highlighting the 40-49 years group as an intermediate group between the young and older adults.

During the VS window, all groups displayed an alpha desynchronization in relation to baseline. In the younger group, the alpha desynchronization was greater in the No-Switch compared to both Switch conditions, but predominantly localised to frontal cortex in the Target Switch condition and left parietal cortex in the No-Target Switch contrast. It could be that this greater alpha desynchronization in frontal and parietal regions reflects increased resources available to attend to the VS in the No-Switch condition compared to the Switch conditions. In addition, the distinct localisations to frontal and parietal cortex could also indicate differences in relation to presence (Target Switch) vs. absence (No-Target Switch), respectively, of a target in the preceding RSVP.

In response to VS presentation, there was a greater alpha decrease in the Target Switch condition compared to the No-Switch condition in the two older groups, whereas the opposite was seen in the younger group. In the RSVP time window that preceded the VS, only the youngest group showed a robust increase in alpha power relative to baseline. It could be that successful inhibition of the RSVP stream in the No-Switch condition is only effectively implemented in the younger group, making more processing resources available to attend to the VS display in the No-Switch condition. In contrast, the middle-aged and older groups may fail to inhibit the irrelevant distractors in the RSVP stream after processing the RSVP Target (which was the first item in the RSVP in the No-Switch condition), and have fewer processing resources available to enhance attention to the VS display. This is further supported by the alpha power decrease in the 60+ group during the RSVP (collapsed cross all conditions, see Figure 7), relative to baseline. However, such an explanation is merely speculative, and future research should aim to thoroughly investigate this hypothesis.

Thus, both theta and alpha signatures revealed widely distributed processing networks in older participants, with a stronger propensity towards frontal involvement compared to the youngest group. However, alpha modulations did not reveal significant correlations with behavioural Switch-Costs, possibly supporting an interpretation in terms of increased neural noise (Shih, 2009; Welford, 1981). However, it should be noted that decreased alpha amplitudes with age might hamper correlational analysis due to a reduction in signal strength. Previous literature has shown that pre-stimulus alpha desynchronization no longer predicts successful stimulus processing in older age (Deiber et al., 2013) as it does in younger adults (Sauseng et al., 2005). The current findings call into question whether pre-stimulus alpha desynchronisation predicts successful target stimulus processing in middle-age. It could be that, in older age, alpha oscillations no longer reliably gate sensory processing (Jensen and Mazaheri, 2010) or enhance attention to visual stimuli (Capotosto et al., 2009; Hanslmayr et al., 2007; Hanslmayr et al., 2005; Klimesch et al., 2007; Rohenkohl and Nobre, 2011; Sauseng et al., 2005; Thut et al., 2006; Yamagishi et al., 2003), possibly placing more demand on top-down attentional control regions. The majority of literature that supports alpha as a sensory gating mechanism has been conducted in younger adults. It is possible that such findings do not generalise to older age groups if processing mechanisms become increasingly altered with age.

The 19-30 years group showed higher Target Switch related theta in parietal regions in comparison to the two older age groups. Posterior parietal activity is usually observed during enhanced attention in young adults (Coull and Nobre, 1998; Li et al., 2013; Madden et al., 2007; Shapiro et al., 2002). However, increased parietal theta in the current task seems to be related to RSVP target processing rather than refocusing attention, as no significant difference in theta was seen between No-Target Switch and No-Switch conditions in the 19-30 years group (Figure 6 panel A). It appears that this parietal theta increase in younger adults reflects enhanced attention directed towards the RSVP target and RSVP target consolidation (Imaruoka et al., 2003). Residual parietal theta in the oldest group was related to reduced Switch-Costs, suggesting that RSVP target processing may influence subsequent switching.

Both the 40-49 and 60+ years groups showed significantly lower occipital and cerebellar theta in Switch conditions (compared to the No-Switch condition), a difference that was not present in the 19-30 years group. Although no significant posterior negative cluster was seen in the 60+ years group in the Target Switch comparison, this could be due to the limited sensitivity of cluster permutation analyses when localising both positive and negative clusters. Indeed, when plotting t-statistics across the entire cortex a similar (non-significant) negative cluster is also visible in the Target Switch condition compared to the No-Switch condition (see Figure SM2). While not surviving robust cluster-based permutation testing theta activity (difference between Target-Switch and No-Switch) in this region revealed a negative correlation with Target RT-Switch-Costs (r=-.44, p=.035) in the 60+ years group.

Weaker posterior theta in the two Switch conditions may be linked to age-related increases in VS RTs in these conditions. Reduced activity in the occipital lobe is consistent with previous findings of age-related reductions in visual cortex activity during visual processing and more generally with the PASA hypothesis (Davis et al., 2008; Huettel et al., 2001; Madden et al., 2002; Ross et al., 1997; Ross et al., 1997).

Higher theta power in the MFG and parietal cortex in the Target-Switch condition (compared to No-Switch) correlated with reduced RT-Switch-Costs in the 60+ years group, implying a supporting role of the MFG (Cabeza et al., 2018; Park and Reuter-Lorenz, 2009; Reuter-Lorenz and Park, 2014). Importantly, the parietal source coordinates were adopted from a theta effect in the youngest group. Thus, it appears that stronger residual parietal theta activity in older individuals, which resembles parietal theta activity in the young group, is beneficial to attentional switching in these older individuals and reflects the *maintenance* of attention mechanisms (Cabeza et al., 2018; Nyberg et al., 2012; Park and Reuter-Lorenz, 2009; Reuter-Lorenz and Park, 2014). This then seems to be complemented by compensatory MFG recruitment in theta. Although the correlation between parietal theta and RT-Switch-Costs did not reach significance due to insufficient power, and thus cannot be regarded as reliable at the current stage, the strength of the correlation (r = −.40) indicates that it is worthy to guide further investigation and future replications.

Based on the observed correlation between MFG theta and reduced Switch-Costs the additional recruitment of frontal regions favours an interpretation in terms of compensatory recruitment of top-down mechanisms of attentional control (Hopfinger et al., 2000) in contrast to an interpretation in terms of de-differentiation (Cabeza, 2002). The additional temporal lobe activity in the 60+ years group on the other hand could indicate further alternative strategies to complete the task, such as episodic memory encoding (Schacter and Wagner, 1999) and/or silent vocalisation (Graves et al., 2007; Hickok and Poeppel, 2007; Hocking and Price, 2009; Smith et al., 1998). A lack of correlation with reduced RT-Switch-Costs means that these theta sources could also reflect de-differentiation. However, an alternative possibility is that they reflect strategies to support task processes unrelated to the speed of switching, such as the maintenance of task goals or storing targets in working memory (e.g., silent vocalisations could facilitate either target recall or the maintenance of task goals). Our observation of increased frontal theta with increasing age contrasts with Cummins and Finnigan’s (2007) findings of reduced theta in frontal EEG electrodes, and instead supports compensatory models of ageing such as STAC (Park and Reuter-Lorenz, 2009; Reuter-Lorenz and Park, 2014) and PASA (Davis et al., 2008). Our findings could therefore have important implications for future work aiming to enhance compensatory recruitment through cognitive training and/or neural stimulation. Ultimately, improving older adults’ attentional flexibility could improve everyday activities such as driving, where one is required to quickly switch between fast changing events in multiple surrounding locations (Huizeling et al., 2020).

## Conclusions

We have replicated the behavioural findings of (Callaghan et al., 2017), observing age-related declines in the ability to switch between temporal and spatial attention. Difficulties in refocusing attention between time and space seem to be accompanied by a deficit in theta power modulation in occipital and cerebellar regions. Older and middle-aged adults’ brains seem to partially attempt to compensate for this posterior theta deficit by recruiting a more extensive frontal network, possibly reflecting increased reliance on top-down attentional control. In addition to more extensive frontal recruitment, the 60+ years group showed recruitment of the temporal lobes, possibly reflecting further attempts of compensation strategies such as episodic memory encoding or silent vocalisation. Efficient (low) Switch-Costs in the youngest group were reflected by parietal theta effects that were absent in both older groups. However, residual parietal theta in the oldest group was related to reduced Switch-Costs, thus, resemblance with efficient processing in the young brain appears to be beneficial for older brains. During the RSVP time window, older adults showed an alpha desynchronization instead of synchronisation, possibly reflecting an enhanced attention strategy rather than an inhibition strategy, as well as a stronger and more extensive task-related alpha power modulation across the cortex. In contrast to theta oscillations, alpha power modulations were not correlated with Switch-Costs, indicating that increases in the extent of power modulation could merely reflect de-differentiation and/or reduced neural precision in older participants.

Overall our results demonstrate that older adults may partially compensate for declines in attentional flexibility with the recruitment of additional theta-related neural mechanisms. These findings have important implications for future work, as they raise the question as to whether this compensatory recruitment can be enhanced with cognitive training programmes. Improving older adults’ attentional flexibility could improve their performance in everyday functions such as driving, where one is required to quickly switch between fast changing events in multiple surrounding locations (e.g. Huizeling et al., 2020).

## Acknowledgements

This research was supported by funding from The Rees Jeffreys Road Fund and by the School of Life and Health Sciences at Aston University. Scanning costs were supported by The Wellcome Trust Lab for MEG Studies and the Dr Hadwen Trust for Humane Research. In addition we would like to thank colleagues at the Aston Brain Centre for assisting with MRI data acquisition.

## Author contributions

Writing – Review & Editing and Resources, E.H., H.W., C.H. and K.K.; Conceptualization and Methodology, E.H., C.H. and K.K.; Formal Analysis, Visualization, Data Collection and Curation, and Project Administration: E.H.; Software, E.H., and H.W.; Writing – Original Draft Preparation, E.H. and K.K.; Supervision and Funding Acquisition C.H. and K.K.;

## Supplementary material

**Figure SM1.**
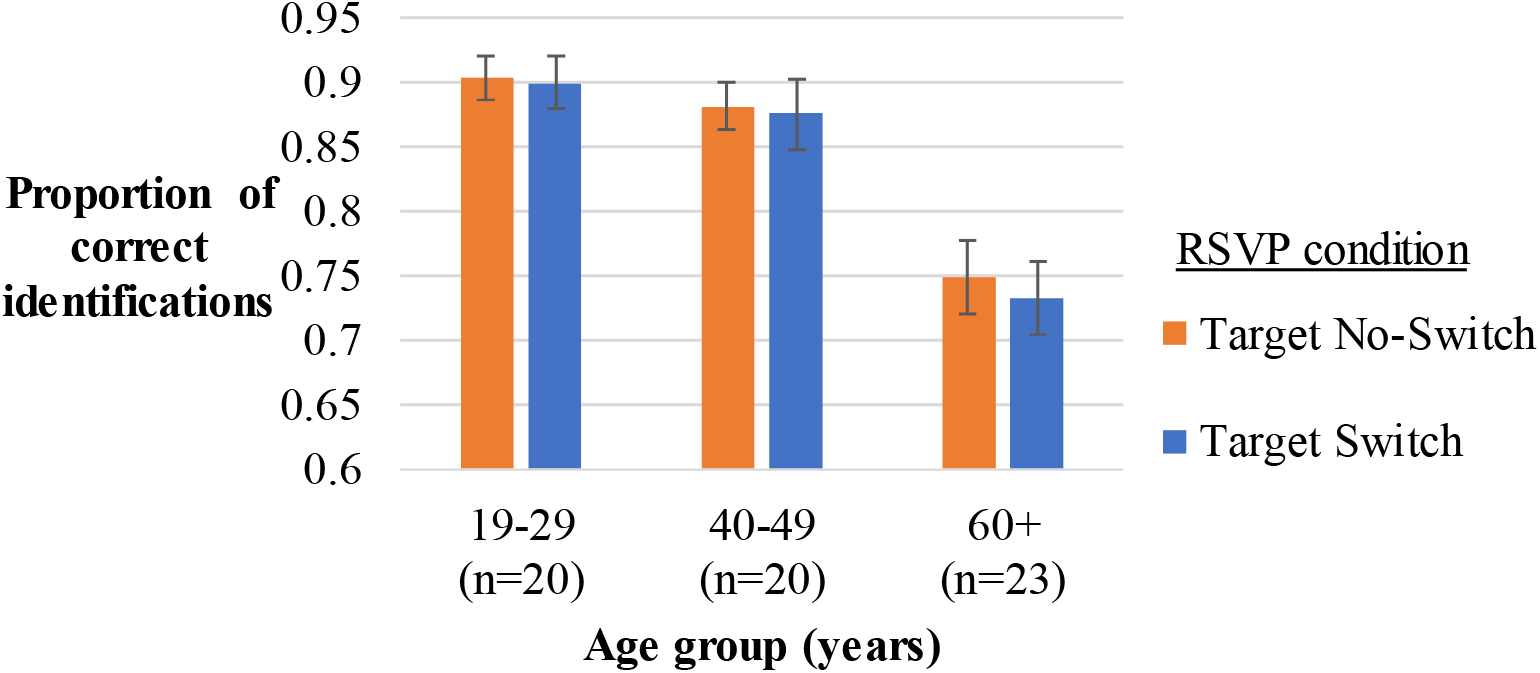
Proportion of correct RSVP target identifications. Vertical bars represent the standard error of the mean.

**Table SM1.**
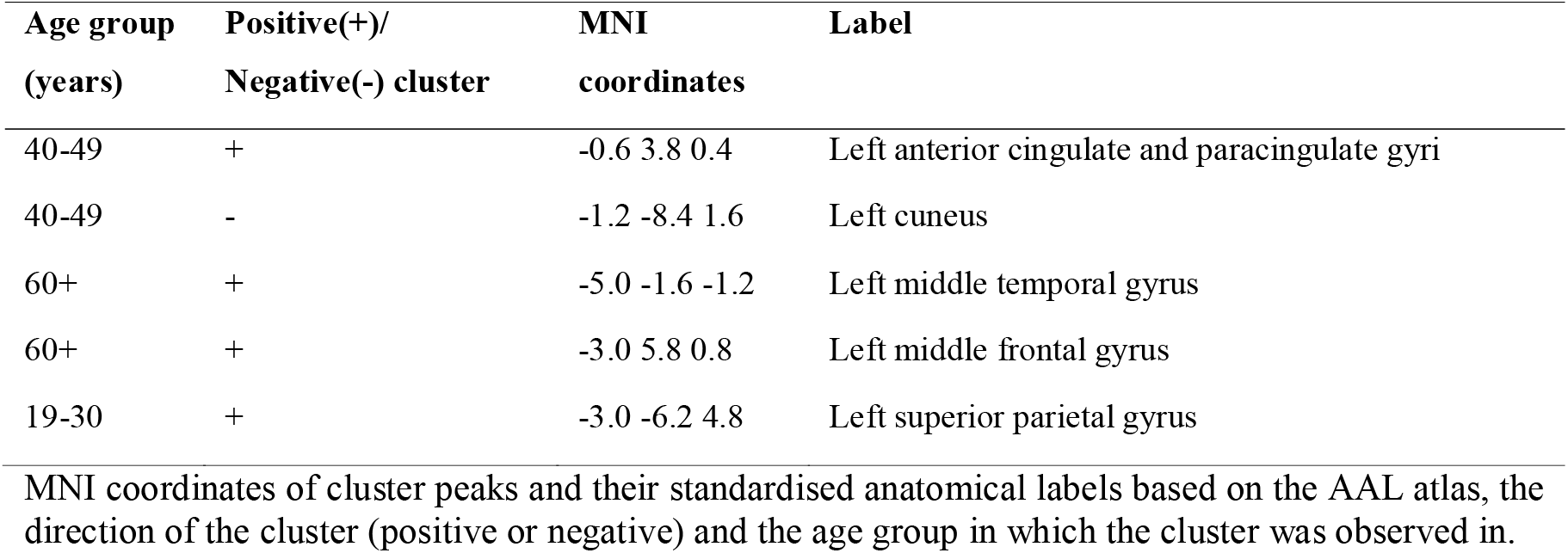
MNI coordinates and atlas labels from which theta power differences between Target Switch and No-Switch conditions were extracted for correlation analyses.

**Table SM2.** MNI coordinates and atlas labels from which alpha power differences between Target Switch and No-Switch conditions during the RSVP time window were extracted for correlation analyses.

| Age group<br>(years) | Positive(+)/<br>Negative(-) cluster | MNI<br>coordinates | Label |
| --- | --- | --- | --- |
| 19-30 | + | 0.2 -2.6 7.8 | Right paracentral lobule |
| 40-49 | + | -1.0 -5.2 4.8 | Left precuneus |
| 60+ | + | -5.4 -4.2 1.8 | Left superior temporal gyrus |

MNI coordinates of cluster peaks and their standardised anatomical labels based on the AAL atlas, the direction of the cluster (positive or negative) and the age group in which the cluster was observed in.

**Table SM3.**
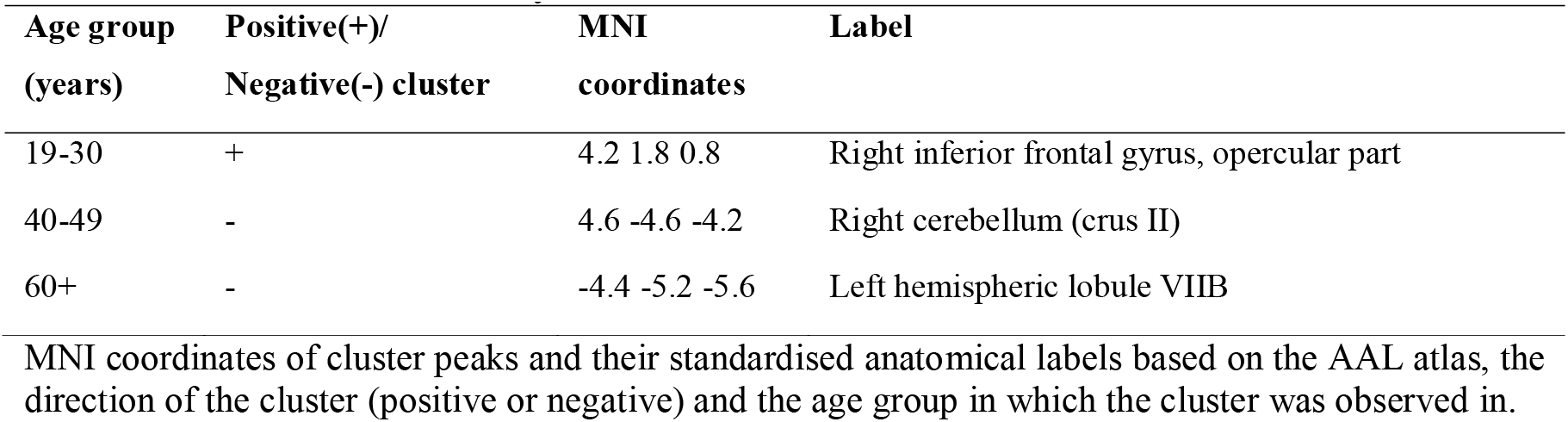
MNI coordinates and atlas labels from which alpha power differences between Target Switch and No-Switch conditions during the VS time window were extracted for correlation analyses.

| Age group<br>(years) | Positive(+)/<br>Negative(-) cluster | MNI<br>coordinates | Label |
| --- | --- | --- | --- |
| 19-30 | + | 4.2 1.8 0.8 | Right inferior frontal gyrus, opercular part |
| 40-49 | - | 4.6 -4.6 -4.2 | Right cerebellum (crus II) |
| 60+ | - | -4.4 -5.2 -5.6 | Left hemispheric lobule VIIB |

MNI coordinates of cluster peaks and their standardised anatomical labels based on the AAL atlas, the direction of the cluster (positive or negative) and the age group in which the cluster was observed in.

**Figure SM2.**
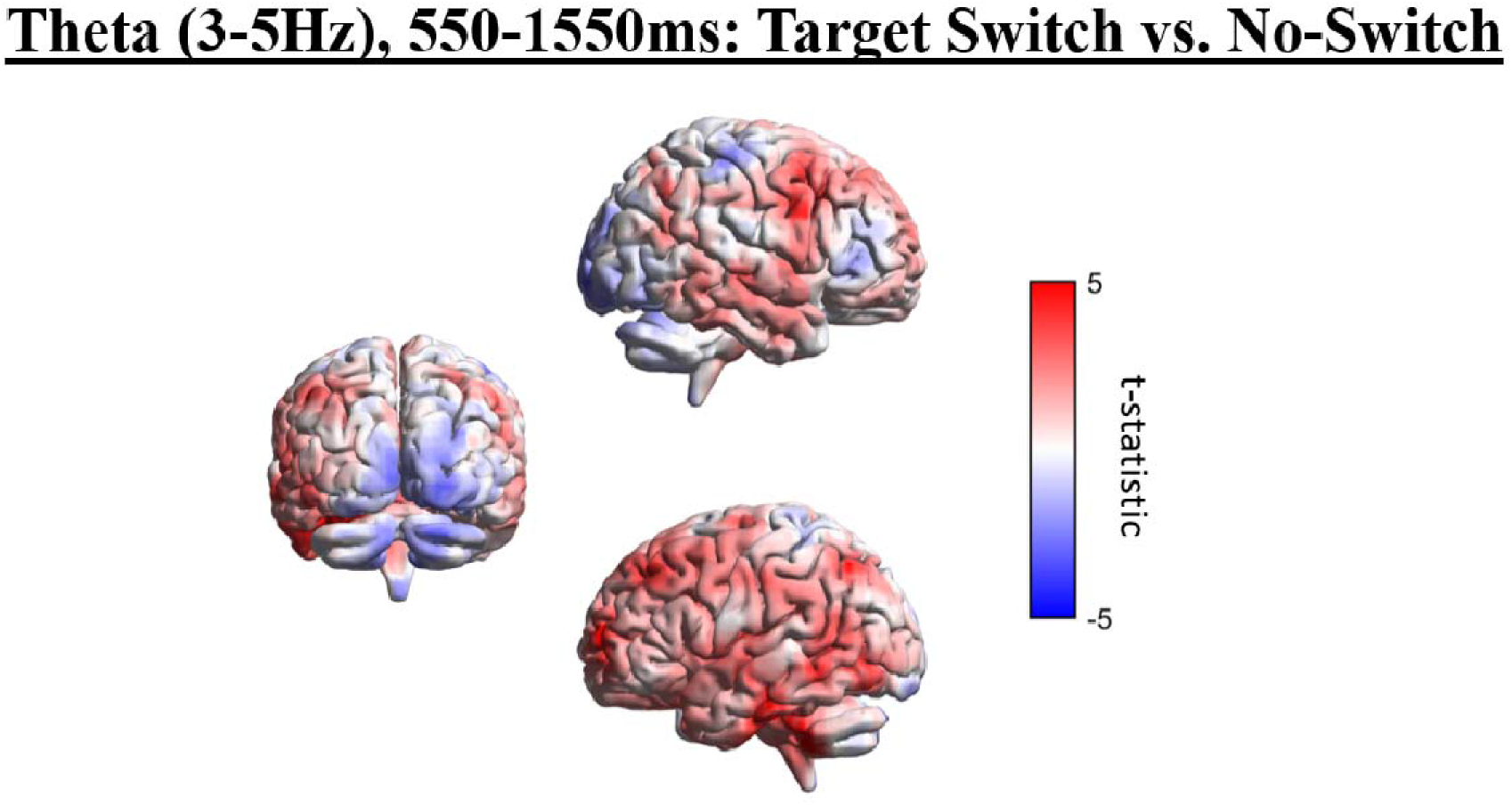
Unthresholded effects in theta (3-5Hz) when contrasting Target Switch and No-Switch conditions in the 60+ years age group. Source plots present *t*-statistics. Note that the most significant positive clusters are identical to the clusters in Fig. 5. This figure illustrates negative effects in the occipital cortex that did not reach significance with the employed robust cluster-based permutation method.

